# Pygopods are an exceptional radiation of snake-like geckos

**DOI:** 10.64898/2026.06.22.733657

**Authors:** Ian G. Brennan, David G. Chapple, J. Scott Keogh, Damien Esquerré

## Abstract

Limb loss in vertebrate animals is surprisingly common despite imposing strong functional constraints. These pressures funnel species towards regions of limited ecological and phenotypic space. To date, snakes have been considered unique in having escaped these limitations. Using a new species-level phylogeny and comparative morphological and dietary datasets, we show that pygopods, a group of limbless Australo-Papuan geckos, have undergone a similar evolutionary trajectory to snakes. Our analyses provide evidence of exceptional morphological and dietary evolution. This is exemplified by strong niche partitioning among genera through dietary specialization and greater than expected dietary disparity. Diversification in pygopods has also been driven by extreme phenotypic evolution, with pygopods encompassing much of the morphological space covered by all other limb-reduced lizards. Interestingly, the diversification of pygopods has resulted in only a modest number of living species, emphasizing the decoupling of diversity and richness possible in adaptive radiations.

## Introduction

Limb reduction and loss is a common trajectory in vertebrate evolution, from birds to amphibians and even mammals. No group, however, has undergone limb loss as extensively or as frequently as squamate reptiles, the group comprising lizards and snakes (Gans, 1975; Lande, 1978; Camaiti et al. 2021). Within squamates, limb reduction has likely evolved more than 50 times and is usually associated with an adaptation to burrowing lifestyles and a transition towards a long snake-like body form (Wiens et al. 2006; Brandley et al. 2008; Camaitti et al. 2021). In many instances, this transition follows a predictable evolutionary path, starting with body elongation, digit loss, then limb length reduction (Gans 1975; Camaiti et al. 2021). This process is best seen in gymnopthalmids (Presch 1975), some skinks (Skinner 2010; Camaiti et al. 2023), and the fossil history of snakes (Rage & Escuillié 2003). The origins of limblessness in other squamate groups remain more enigmatic. One such group is the Pygopodidae, a family of Australo-Papuan geckos that represent one of the most remarkable radiations of limbless reptiles.

Geckos, known for their exceptional ability to climb smooth and even inverted surfaces are perhaps a surprising group to go limbless. There are no known transitional pygopods that might provide insight and there is a tremendous morphological gap between them and their closest fully limbed relatives (Carphodactylidae, Diplodactylidae). Despite this, pygopods have long been associated with geckos (Boulenger 1884), though it took until the molecular era to formalize the position of pygopods within Gekkota (Donnellan et al. 1999). Since then, a number of molecular assessments have verified the placement of pygopods within a Gondwanan-age radiation of Australian geckos that includes the Diplodactylidae and Carphodactylidae (Han et al. 2004; Oliver & Sanders 2009; Gamble et al. 2011; Skipwith et al. 2019). What has evaded us though, is an accurate understanding of the relationships among pygopods. Deeper relationships within the family have been hard to resolve partly due to extreme morphological divergence and ecological differences among genera (Jennings 2021).

The tremendous breadth in ecologies and morphologies of pygopods is largely what makes them remarkable. Together, they span large (750 mm) nocturnal lizard-eating ambush predators like *Lialis* to small (100 mm) diurnal and fossorial ant-eaters like *Aprasia*. However, they are not broadly distributed (Australo-Papuan endemics) or particularly species rich (∼50 species). But the contrast of their low richness and high ecomorphological diversity has made them a classic example of the decoupling of speciation and divergence that is possible in adaptive radiations (Losos & Mahler 2010; Esquerre et al. 2022). This is of interest because most other limbless squamates show narrow ecological breadth and are largely derivations on the same fossorial or cryptozoic morphologies (Wiens et al. 2006; Anelli et al. 2024). One notable exception is snakes. Through an incredible adaptive display, snakes have radiated into an array of varieties found below ground, high in the trees, and even in the middle of the ocean (Title et al. 2024). To emphasize the novelty of pygopods we use morphological and dietary data to compare them to other squamate radiations, particularly snakes, which are commonly viewed as evolutionarily singular and unique.

Here we suggest that the transition towards limblessness in squamate reptiles imposes ecomorphological constraints. While snakes have no doubt escaped this generalisation, it is unclear if pygopods show greater ecological and morphological diversity than expected given their age. To address this we investigated the macroevolution of pygopods using a sequence-capture dataset to reconstruct the phylogenetic relationships among all described species and morphological and diet data to explore their diversification. Our findings highlight the importance of niche partitioning in a radiation of limited species richness.

## Materials and Methods

### Molecular Data Collection and Phylogenetic Analyses

We assembled a sequence-capture dataset from 160 pygopods representing all 50 recognized species and relevant outgroups (Table S1). For molecular sampling we used the Squamate Conserved Loci (SqCL) kit (Singhal et al. 2017), which comprises ∼5,000 ultra-conserved elements (UCEs), ∼400 anchored hybrid enrichment loci (AHEs), and ∼40 legacy genes commonly used in squamate molecular phylogenetics. Sample collection and preparation were completed as part of the Australian Amphibian and Reptile Genomics initiative (AusARG) following the protocol outlined in Tiatragul et al. (2023). We processed raw sequence data using the *pipesnake* workflow (Brennan et al. 2024), including estimating individual genealogies using IQ-TREE2 (Minh et al. 2020) and species trees using the coalescent-consistent methods wASTRAL-hybrid and ASTRAL-IV (Zhang & Mirarab 2022). Additional details regarding molecular data are presented in the Supplement.

### Divergence Dating

To estimate divergence times among taxa we applied a series of fossil and secondary calibrations in MCMCTree (Rannala & Yang 2007) as outlined in the Supplement and Table S2. We started by trimming our wASTRAL tree down to a single representative for each ingroup species. We reduced our molecular data to exonic markers in the AHE loci, divided the loci into four rate-based partitions, and retained only the first and second codon positions. For each partition we estimated the approximate likelihoods and branch lengths (usedata = 3), and ran four replicate MCMCTree analyses on the concatenated output (usedate = 2). We then combined the mcmc files to summarize divergence times on our tree.

### Morphological and Dietary Evolution

To investigate patterns of tail evolution across squamates we generated a dataset by measuring museum specimens and collecting data from the literature. We focused on two traits commonly recorded for squamate reptiles, snout–vent length, and tail length. Together these traits are generally indicative of total animal size, but differ in their relative proportions across squamates. Typically species fall into one of two categories: (1) long bodied and short tailed as in snakes, amphisbaenians, and dibamids, or (2) short bodied and long tailed as in most fully limbed and limb-reduced lizards. Our dataset includes more than 11,500 individuals across 3,105 species and 80 squamate families. This builds on extensive datasets compiled by Jennings (2002), Brandley et al. (2008), Meiri (2024), Title et al. (2024), and Anelli et al. (2024). To visualize trends in relative tail lengths we started by plotting the ratio of tail length (TL) to snout–vent length as a function of total length (TL/SVL ∼ log(Total)) (Fig.2). We then fit separate linear models to each family (Fig.S4). We also plotted these patterns across major morphological groups by splitting squamates into pygopods, fully limbed lizards, limb-reduced lizards, and snakes, following the classifications of SquamBase (Meiri 2024).

To better understand the tempo and mode of elongation through tail evolution we undertook a trait modelling exercise. We started by grafting our dated pygopod tree onto the squamate topology of Title et al. (2024) and we retained squamate taxa that were available in both the tree and morphological dataset (TL/SVL). Using these data we fit a series of models ranging from an unbiased random walk to rate-variable and trended. The most basic model—Brownian Motion—allows phenotypes to evolve through incremental random change, but at deep timescales and when traits may show evolutionary rate heterogeneity this model is expected to perform poorly. The most complex—BayesTraits V4’s fabric model (Pagel et al. 2022)— also follows a process of gradual change, but allows rates to shift at discrete points (changes in *evolvability, v*) on the tree and allows traits to change rapidly along individual branches through directed evolution (*trends*, β). We fit these models and others (Early Burst, single-peak Ornstein-Uhlenbeck) in BayesTraits V4 and estimated marginal likelihoods using a stepping-stone sampler to compare model fit. More details on our modelling exercise can be found in the Supplement.

To investigate patterns of diet breadth evolution across squamates we built on existing datasets (Title et al. 2024; Cavalcanti et al. 2025) by collecting data from the literature. Our dataset includes diet records for more than 90,000 individuals across 1,738 species and 63 squamate families. This new dataset significantly expands on efforts elsewhere by incorporating nearly 200 additional species, primarily lizards, including many limb-reduced species (e.g. *Dibamus*) and those with dietary items not previously captured (e.g. seaweed eating *Amblyrhynchus cristatus*, and frugivorous lizards like *Conolophus pallidus* and *Varanus mabitang*). To process and analyze the data we followed the methodology of Title et al. (2024) in standardizing diet proportions to make lizards and snakes comparable, and the analytical framework of Grundler & Rabosky (2021). Briefly, this framework calculates a phylogenetically informed estimate of dietary niche for each species by discretizing the multivariate diet space (32 dimensions) into *K* niche states, then samples from the diet variables using MCMC. For the sake of visualizing this diet space we used principle components analysis for dimensionality reduction. To compare dietary disparity among squamate families we calculated the mean pairwise distance in euclidean space among all species in each family and carried out pairwise family comparisons with species permutations to assess significance. To determine how dietary disparity compares to a null diffusion process we estimated evolutionary rates for each dietary category under Bounded Brownian Motion, then simulated 500 datasets using estimated parameters (Fig.S9).

## Results

### Phylogenomics and Divergence Dating

We present a new phylogenetic hypothesis of relationships for all 50 species of pygopodid geckos based on sequence capture data. We recovered 5406 loci covering 6.3 mbp with a mean locus alignment length of 1258 bp (max. = 2508, min. = 479) and sample occupancy of 133 individuals (± 24).

Coalescent-consistent analyses using wASTRAL-hybrid and ASTRAL-IV provided topologies that are well resolved with high support (>90 local posterior probability) across nearly all branches of the tree, including good resolution among genera (Fig.S1,S2,S3). Phylogenetic uncertainty is limited to situations of intraspecific sampling. Internal branches separating some genera are short and show moderate levels of incomplete lineage sorting, resulting in lower gene concordance factors (Fig. S4). This is notable in the splits among genera to the exclusion of *Delma*. An exciting finding places the fossorial *Ophidiocephalus* and arboreal *Pletholax* as sister genera, highlighting the ecological and phenotypic evolution of pygopod diversity.

Pygopods diverged from carphodactylids in the Early Eocene (∼50 ma), and we reconstruct a long pygopodid stem lineage leading to a crown divergence of living species in the Late Oligocene (∼28 ma) (Fig.S5). A series of rapid splits deep in the pygopodid tree roughly around the Oligocene Miocene transition (∼23 ma) separate most major clades. The majority of extant diversity accumulated during the Middle to Late Miocene (13-5 ma) driven by the most species rich genus *Delma*.

### Morphology

Allometric trends in tail-to-body length vary among and within families and morphological groups (limbed, limb-reduced, snake, pygopods) (Fig.2). Snakes show a highly cohesive shallow slope (B = 7.2e-5, SE = 6.6e-6), with limited variation in tail/body ratios (Table S3). This is most extreme in the fossorial families Leptotyphlopidae, Typhlopidae, and Uropeltidae, (tail-to-body ratios <0.01) and this trend is shared with other fossorial squamate groups such as amphisbaenians (Amphisbaenidae, Blanidae, etc.) and dibamids (Table S4, Fig.S6). Lizard groups with varied ecologies such as the Agamidae, Cordylidae, Gymnophthalmidae, Pygopodidae, Scincidae, and Varanidae show a greater spread of points. The Pygopodidae show several distinct trajectories corresponding to the fossorial genus *Aprasia* and terrestrial relatives in *Delma* and *Pygopus*. Species with extreme tail-to-body ratios (>3.5) include both limb-reduced species such as the pygopodids *Delma concinna* and *Delma labialis*, and the cordylids *Chamaesaura tenuior* and *Chamaesaura miopropus*, as well as fully-limbed species such as the lacertids *Takydromus sexlineatus* and *Takydromus sauteri*, and the agamids *Diporiphora superba* and *Hypsilurus longii*.

The best fitting evolutionary model was the *fabric* model (Table S3), which identified a complex interplay of rate heterogeneity and trended changes in tail evolution across squamates (Fig.2, S7). Notably there are extreme increases in evolutionary rate and phenotype in pygopodids, particularly in *Delma* (β = 1.67; *v* = 28.58), *Pletholax* (β = 1.76), and *Lialis/Pygopus* (β = 0.51). These occur among a number of other rate and mean phenotype changes across squamates. In snakes there is a dramatic trait decrease at the base of snakes (β = -1.35), and significant rate decreases in Cylindrophiidae/Uropeltidae (*v* = 0.02) and Typhlopidae (*v* = 0.22).

### Diet

We retained 31 families with at least five species with diet data for investigating dietary disparity. Among squamates, pygopodids (50 species) show one of the greatest disparities in diets, third only to varanid lizards (88 species) and colubrid snakes (2192 species), two groups known for their extreme diversity (Fig.2, Table S6). Differences in breadth among these groups are not significant (Fig.S8). Thirteen families showed pairwise distances that were consistent with expectations under a neutral dietary diffusion process (Fig.S9). Fourteen families showed lower than expected mean pairwise distances indicating narrower dietary divergence among species. Eight families showed greater than expected mean pairwise distances indicating significant dietary disparity. Of the eight families with wider dietary disparities, four are snakes (Colubridae, Elapidae, Homalopsidae, Viperidae) and four are lizards (Agamidae, Liolaemidae, Pygopodidae, Varanidae).

## Discussion

Limb reduction is a common evolutionary trend in squamate reptiles. In lizards, limb loss is typically associated with a fossorial lifestyle and relatively predictable morphology. Here, we provide evidence that pygopod geckos—similar to snakes—have escaped these trends and show notable ecomorphological divergence.

### Phylogenetics and Diversification

Pygopods are absolute outliers in the world of geckos. They are ecologically, behaviorally, and phenotypically divergent from this group, and superficially show greater similarity with snakes. Adaptive diversification into highly modified and differentiated genera has presented unique challenges in understanding their evolution. Traditional morphological methods drew generalist species (*Delma, Pygopus*) together towards the base of the tree (Kluge 1976; Conrad 2008; Pyron 2017). While molecular data are insensitive to these biases, the rapid early diversification of pygopods continued to hamper small mito-nuclear datasets, providing little intergeneric resolution (Jennings et al. 2003; Brennan et al. 2016). Even modern genome-wide datasets (Skipwith et al. 2019) have failed to provide a reliable tree of life for these unique animals (see Jennings 2021 for extensive coverage).

Here, we present a well-resolved, fully sampled phylogenomic hypothesis of the Pygopodidae, with a novel intergeneric configuration and a few surprises. We support past assessments that placed *Delma* as sister to the remaining genera (Jennings et al. 2003; Brennan et al. 2016; Skipwith et al. 2019; Jennings 2021), and also support Skipwith et al. (2019) in identifying a clade of large-bodied genera in *Lialis, Paradelma*, and *Pygopus*. Our most surprising finding is the sister relationship between *Ophidiocephalus* and *Pletholax*. These two genera are currently separated by more than 20 million years and 1500 kilometers across outback Australia. Their dramatically different ecologies (fossorial vs. shrub swimming) indicates repeated and rapid shifts in habitat use and foraging strategy possibly in response to changing Miocene habitats (Jennings et al. 2003). This suggests ecological lability was possibly a trademark of early pygopods. However, extant diversity is comparatively young (stem age ∼50 my, crown age ∼25 my) and likely the result of post-extinction recovery (Brennan & Oliver 2017). Begging the question, what were early pygopods like?

### Radiation through Tail Evolution

In the squamate body, elongation is the primary axis for morphological change and occurs most commonly by adjusting the body-to-tail length ratio. In non-snake squamates, limblessness is typically associated with a narrow region of ecomorphospace characterised by long-bodied, short-tailed fossorial species (Wiens et al. 2008; Brandley et al. 2008; Anelli et al. 2024). This morphotype is found across amphisbaenians, dibamids, gymnophthalmids, anniellids, and various groups of skinks, as well as the pygopod genera *Aprasia* and *Ophidiocephalus*. Pygopods have escaped this motif by dramatically varying their tail lengths from less than one-third (*Aprasia*) to more than four times (*Delma*) their body length. Our modelling indicates this has happened through both increases in evolutionary rates and selection, possibly driven by ecology. However, pygopods are not the only group to expand the morphospace through tail extension. Other long-tailed limb-reduced squamates such as anguids (*Ophisaurus*), cordylids (*Chamaesaura*), and gerrhosaurids (*Tetradactylus*) have followed a similar path. Though intriguing, these groups are species poor and ecologically uniform. The morphological diversity of these additional groups is captured entirely within the radiation of pygopods. Grass or shrub-swimming *Chamaesaura* and *Tetradactylus* are ecologically similar to *Pletholax* and some *Delma* (Berger-Dell’mour 1985; Du Toit 2001).

Importantly, pygopods demonstrate that elongation in limbless squamates need not follow the highly conserved “snake model” of body elongation coupled with tail reduction (Fig.2). Instead, diversification along the tail axis appears to be the key driver of pygopod morphological disparity. This reflects differences in locomotion, habitat use, and predator avoidance. In some *Delma* species, the tail is used for propulsion, lifting the animal clear from the ground in successive saltations (Bauer 1986). In all pygopods tail autotomy is common (Jennings 2002) suggesting it is an effective tool for escape. This practice is emphasized in some *Aprasia* which have short tails that are marked conspicuously like the head, possibly to confuse predators (such as *Aprasia smithi*). The variety of strategies in pygopods emphasizes the decoupling of elongation pathways and challenges the prevailing view that limblessness imposes strong constraints on body form. Instead, it suggests that alternative developmental and functional routes can generate substantial morphological diversity.

### Dietary Specialization and Niche Partitioning

Patterns of dietary partitioning and evolution in pygopods further reinforce their exceptional nature. While most non-snake limb-reduced squamates display generalist insectivorous diets (see data in Title et al. 2024 and Cavalcanti et al. 2025), our results show that pygopod species have comparatively narrow diets that are more disparate than expected under a null evolutionary model. This partitioning may explain the sympatric coexistence of highly specialized and generalist lineages (Losos & Mahler 2010). Along the coast of Western Australia, six or more species of five pygopod genera may exist in sympatry (Atlas of Living Australia 2026). In these coastal habitats specialists like lizard-eating *Lialis* and ant-eating *Aprasia* define major axes of dietary variation and live alongside species with broader dietary breadths like *Delma* and *Pygopus*. Overdispersion in pygopod prey makes for one of the most varied dietary compositions of any squamate family (Table S6, Fig.S8), comparable to that of major snake groups like colubrids and elapids. However, dietary differences alone do not explain this level of species richness.

Like all good adaptive radiations, ecological partitioning has likely facilitated pygopod richness and community assembly. In addition to dietary diversification, pygopods have also divided the diel cycle and microhabitats (Patchell & Shine 1986; Gurgis et al. 2021). As a result, a diurnal fossorial species like *Aprasia repens* likely never encounters a diurnal arboreal species like *Pletholax gracilis*, a crepuscular terrestrial species like *Pygopus lepidopodus*, or a nocturnal terrestrial species like *Lialis*, even though all these species live in the same habitat. This level of partitioning is extreme and demonstrates that high ecomorphological disparity can evolve in relatively small clades. Diversification across multiple axes (morphology, diet, habitat) underscores the exceptional nature of the pygopod radiation.

## Acknowledgements

This project represents an output of the Australian Amphibian and Reptile Genomics initiative (AusARG) with data generation generously funded by BioPlatforms Australia and salary support through an Australian Research Council Discovery Grant (DP210100820 to JSK; LP170100012 and FT200100108 to DGC). We appreciate the provision of computing and data resources provided by the Australian BioCommons Leadership Share (ABLeS) program and Seqera Tower service. These programs are co-funded by Bioplatforms Australia (enabled by NCRIS), the National Computational Infrastructure and Pawsey Supercomputing Centre. Special thanks to Sophie Mazard, Ziad al Bkhetan, Wes Read, and Sarin ‘Putter’ Tiatragul. We thank our many Australian partner institutions (Australian Museum, Museum and Art Gallery of the Northern Territory, Queensland Museum, Museums Victoria, South Australian Museum, Western Australian Museum) and their curators and collections managers, who made this work possible through generous tissue loans and collections access. Sincere thanks to W. Bryan Jennings and Aaron Bauer for their roles in developing and supporting IGB’s continued interest in pygopods. We declare no conflicts of interest.

## Figure Legends

**Figure 1.**
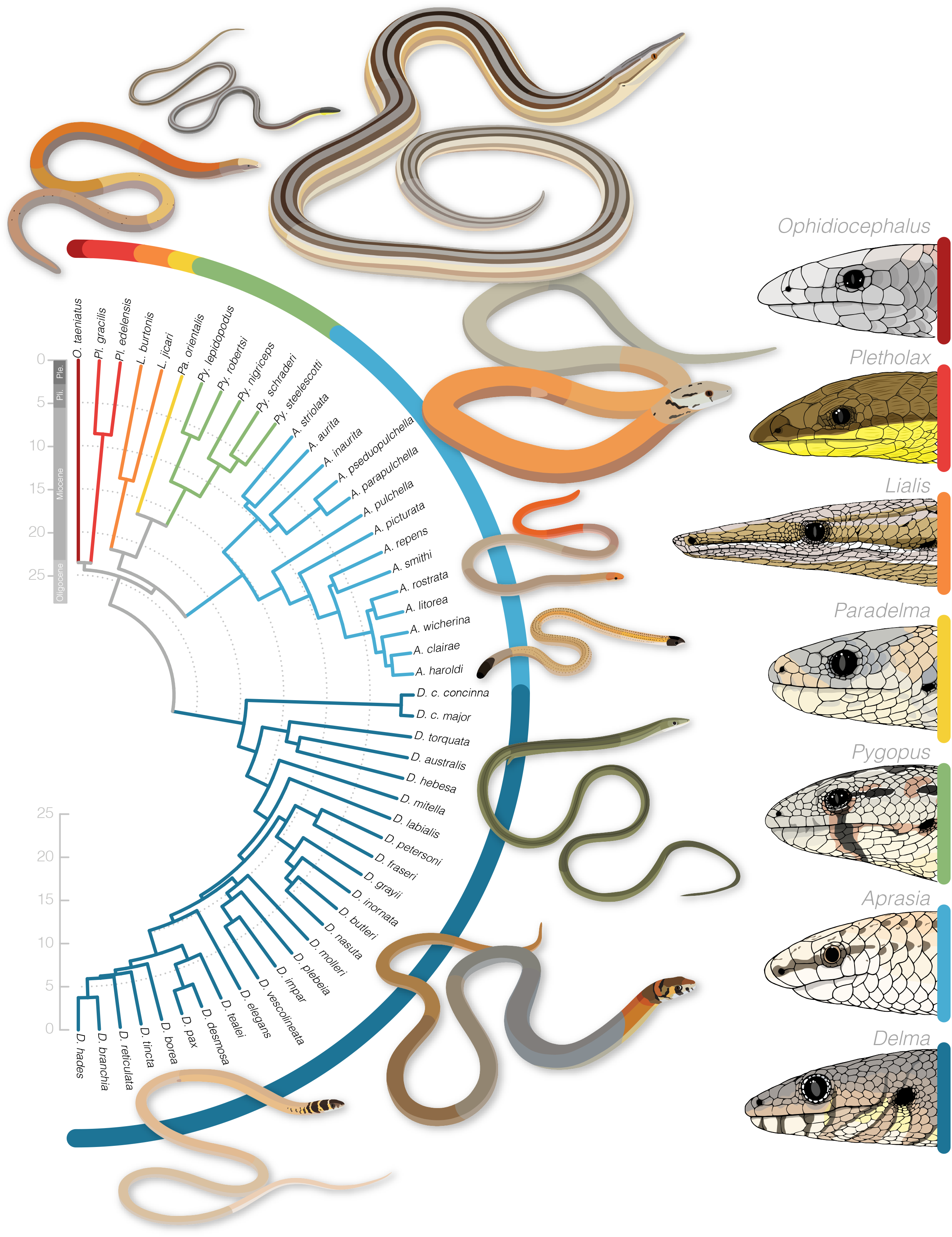
The diversification of living pygopods began around the Oligocene-Miocene boundary and rapidly diverged into the major ecomorphs which largely correspond to genera. ASTRAL species tree time-calibrated with MCMCTree. Diversity of the group is roughly split between the primarily generalist terrestrial *Delma* and remaining more specialized genera. Illustrations surrounding the tree and of heads at far right highlight the morphological diversity of the group, from stout ambush predators to gracile grass swimmers. Full-body illustrations by Wes Read, clockwise from top are *Ophidiocephalus taeniatus, Pletholax gracilis, Lialis burtonis, Pygopus lepidopodus, Pygopus lepidopodus, Aprasia inaurita, Aprasia smithi, Delma concinna, Delma petersoni*, and *Delma branchia*. Illustrations are available from github.com/IanGBrennan/Figure_Critters.

**Figure 2.**
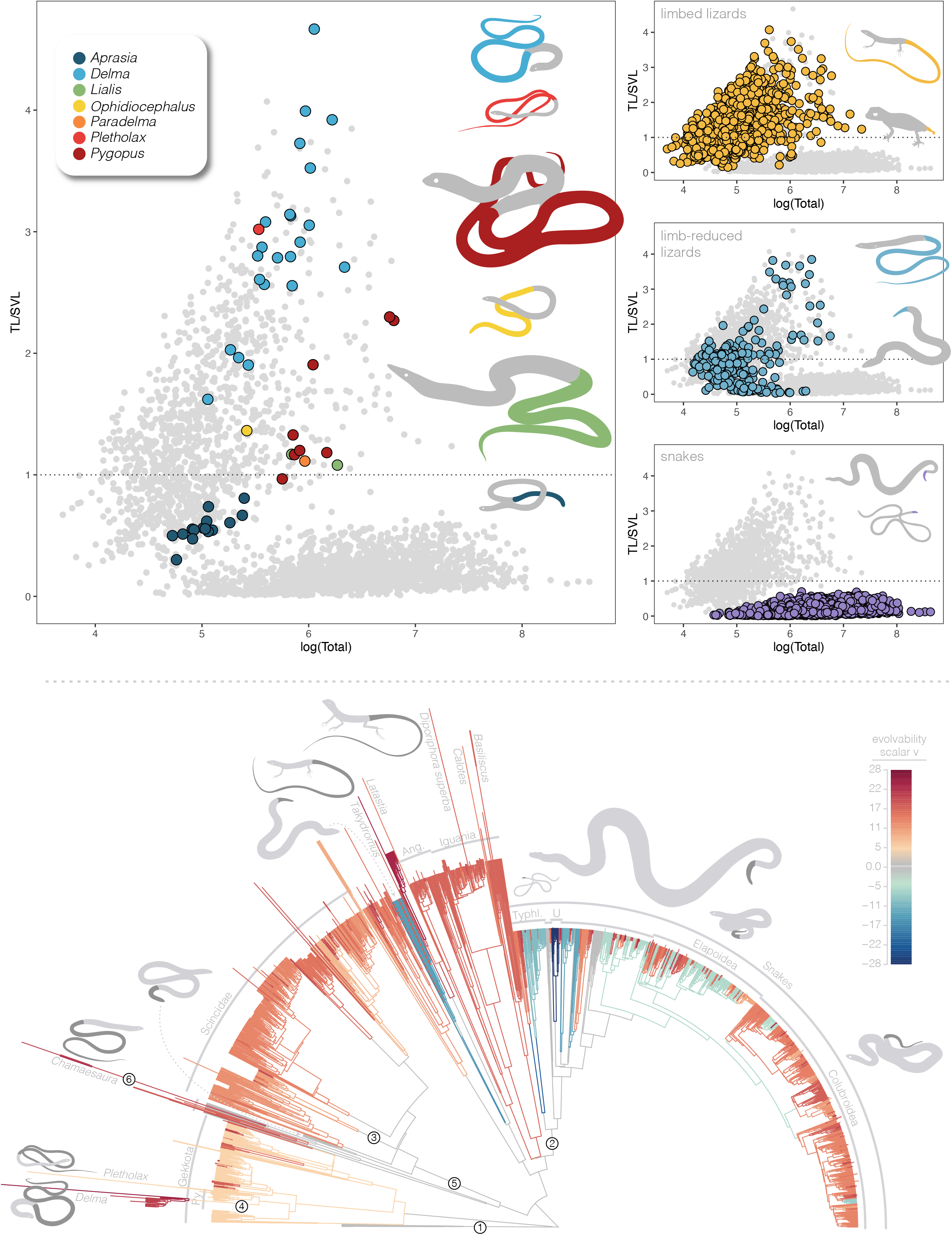
Pygopods cover nearly the entire spectrum of squamate diversity in relative tail-to-body length. The large plot at top-left shows the allometric trend in tail-to-body ratio and with pygopod species colored by genus, compared to all squamates shown in smaller grey points in the background. The dotted line indicates a 1:1 relationship where body length and tail length are equivalent. At top-right, plots from top to bottom show the distribution of these morphologies across fully limbed (top) and limb-reduced lizards (middle), and snakes (bottom). The relative invariance of body proportions in snakes shown by the flat distribution of points along the x axis contrasts with the variety of forms found in other lizard groups. The tree below highlights the varied history of tail evolution across squamates as estimated under the fabric model. Branches are colored according to relative evolutionary rate scalar *v* (warm colors are faster than the background rate, cool colors are slower, grey is background). Branch lengths are scaled to reflect directional phenotypic trends β. Shortened branches like at the base of dibamids (1), snakes (2), and the scincid genus *Brachymeles* (3) indicate directional trends towards shorter relative tail-to-body lengths. Lengthened branches like at the base of the pygopod genus *Delma* (4), the Scincidae (5), and the cordylid genus *Chamaesaura* (6) indicate directional trends towards longer relative tail-to-body lengths.

**Figure 3.**
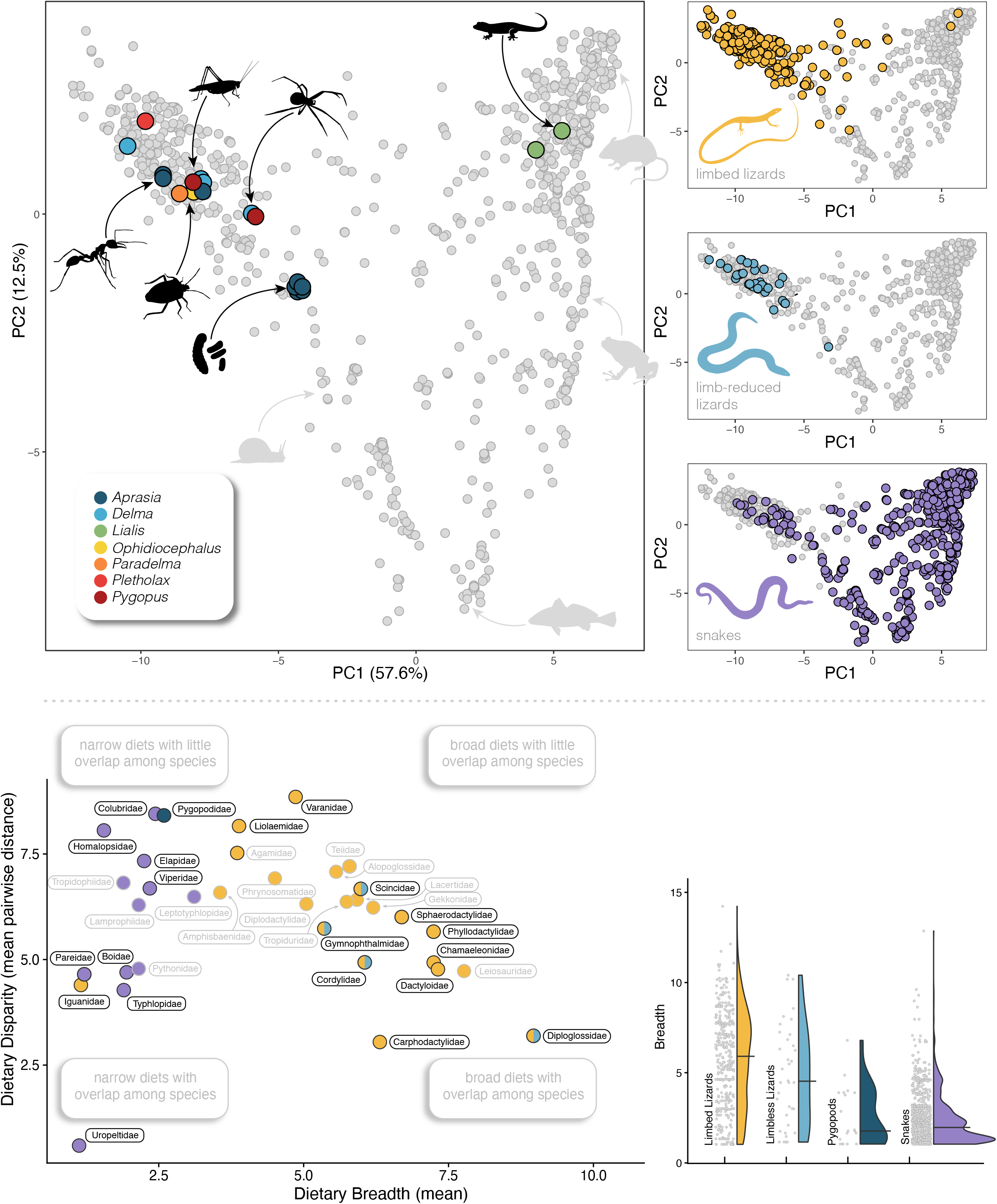
As a group, pygopods show exceptional dietary diversity driven by narrow breadths of individual species. Prey specialization in *Lialis* (squamates) and *Aprasia* (ants, their larvae and eggs) captures two of the major dietary axes. Remaining genera are primarily generalists with some dietary preferences such as an affinity for spiders in *Pygopus*. The plot at upper left shows the distribution of pygopod species, colored by genus, compared to all squamates shown in smaller grey points in the background. Major dietary preferences are illustrated with black silhouettes and include squamates, orthopterans, spiders, hymenopterans, hemipterans, and insect larvae and pupae. At right, plots from top to bottom show the distribution of diets across fully limbed (top) and limb reduced lizards (middle), and snakes (bottom), illustrations not to scale. As demonstrated by Title et al. (2024) snakes show incredible dietary breadth and specialisation that encompasses nearly all lizard diets, except herbivory. Lower plots, pygopods show dietary disparity greater than other limb reduced squamates combined (save snakes) and also show high levels of specialisation (low breadth).

