## Supplement for "Pygopods are an exceptional radiation of snake-like geckos"

**Supplementary References** **25**

### Supplementary Information

All data, scripts, figures, and files necessary to replicate this work are available on the project GitHub page IanGBrennan/Pygopodidae

#### Molecular Sampling and Analysis

##### Taxon Sampling

We assembled a sequence-capture dataset from 171 pygopods representing all 50 recognized species (Table S1). We included outgroup representatives from diplodactylid (*Oedura luritja*) and carphodactylid (*Saltuarius cornutus*) geckos and used tuatara (*Sphenodon punctatus*) to root the tree.

##### Molecular Data Collection and Processing

Molecular sampling implemented the Squamate Conserved Loci (SqCL) kit (Singhal et al. 2017), which comprises ~5,000 ultra-conserved elements (UCEs), ~400 anchored hybrid enrichment loci (AHEs), and ~40 legacy genes commonly used in squamate molecular phylogenetics. Sample collection and laboratory preparation, including extraction, shearing, size selection, library generation, and hybridization capture were completed as part of the Australian Amphibian and Reptile Genomics initiative (AusARG) under the Phylogenomics Working Group following the protocol outlined in Tiatragul et al. (2023).

Raw sequence reads are available from the Bioplatforms Australia Data Portal. We processed raw sequence data using the *pipesnake* workflow (Brennan et al. 2024). A list of all software and versions used is included in Supplement/software_versions.yml. In brief, the pipeline concatenates individual read files, then removes duplicate reads via BBMAP (Bushnell 2014), trims adapters and barcodes with TRIMMOMATIC (Bolger et al. 2014), identifies read pairs with PEAR (Zhang et al. 2014), removes off-target reads with BBMAP, assembles reads to contigs with SPAdes (Prjibelski et al. 2020), maps contigs to targets with BLAT (Kent 2002) and extracts the best hit for each target, carries out preliminary alignments with MAFFT (Katoh & Standley 2013), before refining the alignments with ClipKit (Steenwyk et al.), estimates locus trees with IQ-TREE2 (Minh et al. 2020), and a species tree with weighted ASTRAL hybrid (Zhang & Mirarab 2022). Alignments were further manipulated using SEGUL (Handika & Esselstyn 2024). *pipesnake* is a highly reproducible workflow that provides consistent results with little user input.

**Molecular Summary Statistics** Summaries of molecular sampling are available in Alignments/Alignment_Summary/ on a per-locus and per-taxon basis. Briefly, there are **5,406** loci covering **7,070,483** aligned sites across **926** million characters. Average locus length is **1,307** bp (min. = 437, max. = 3,514) with an average sample occupancy of **130** individuals per alignment.

A CSV file of this project's molecular sampling information is available at Sampling/Pygopodidae_SampleInfo.csv

Table S1: Table of molecular samples. Fields indicate the number of loci recovered per sample, as well as the project under which the data were generated.

| lineage | AHE | gene | uce | total_loci | Targets | Source |
|---|---|---|---|---|---|---|
| Carphodactylidae_Saltuarius_cornutus_A002601 | 355 | 32 | 4815 | 5202 | SqCL | AusARG (this project) |
| Diplodactylidae_Oedura_luritja_CCM5974 | 354 | 29 | 4827 | 5210 | SqCL | AusARG (this project) |
| Pygopodidae_Aprasia_aurita_ABTC57777 | 294 | 28 | 4056 | 4378 | SqCL | AusARG (this project) |
| Pygopodidae_Aprasia_aurita_ABTC58595 | 320 | 29 | 4365 | 4714 | SqCL | AusARG (this project) |
| Pygopodidae_Aprasia_clairae_ABTC132623 | 235 | 25 | 3345 | 3605 | SqCL | AusARG (this project) |
| Pygopodidae_Aprasia_clairae_ABTC132624 | 328 | 27 | 4222 | 4577 | SqCL | AusARG (this project) |
| Pygopodidae_Aprasia_haroldi_WAM_TR1528 | 258 | 28 | 3752 | 4038 | SqCL | AusARG (this project) |
| Pygopodidae_Aprasia_inaurita_ABTC95771 | 325 | 29 | 4297 | 4651 | SqCL | AusARG (this project) |
| Pygopodidae_Aprasia_inaurita_WAMR137756 | 268 | 28 | 3583 | 3879 | SqCL | AusARG (this project) |
| Pygopodidae_Aprasia_litorea_WAMR141606 | 324 | 26 | 4376 | 4726 | SqCL | AusARG (this project) |
| Pygopodidae_Aprasia_litorea_WAMR151306 | 254 | 29 | 3901 | 4184 | SqCL | AusARG (this project) |
| Pygopodidae_Aprasia_parapulchella_AMSR127439 | 351 | 30 | 4714 | 5095 | SqCL | AusARG (this project) |
| Pygopodidae_Aprasia_picturata_WAMR126998 | 318 | 29 | 4290 | 4637 | SqCL | AusARG (this project) |
| Pygopodidae_Aprasia_picturata_WAMR131647 | 302 | 28 | 4209 | 4539 | SqCL | AusARG (this project) |
| Pygopodidae_Aprasia_pseduopulchella_ABTC70373 | 305 | 26 | 4154 | 4485 | SqCL | AusARG (this project) |
| Pygopodidae_Aprasia_pseudopulchella_SAMAR46279 | 352 | 31 | 4742 | 5125 | SqCL | Title & Singhal et al. 2024 |
| Pygopodidae_Aprasia_pulchella_163481 | 19 | 0 | 4586 | 4605 | UCE | Skipwith et al. 2019 |
| Pygopodidae_Aprasia_pulchella_WAMR119804 | 271 | 28 | 3950 | 4249 | SqCL | AusARG (this project) |
| Pygopodidae_Aprasia_pulchella_WAMR153937 | 254 | 29 | 3499 | 3782 | SqCL | AusARG (this project) |
| Pygopodidae_Aprasia_repens_172504 | 17 | 0 | 4399 | 4416 | UCE | Skipwith et al. 2019 |
| Pygopodidae_Aprasia_repens_WAMR119421 | 302 | 28 | 4153 | 4483 | SqCL | AusARG (this project) |
| Pygopodidae_Aprasia_repens_WAMR127404 | 312 | 28 | 4268 | 4608 | SqCL | AusARG (this project) |
| Pygopodidae_Aprasia_repens_WAMR129543 | 310 | 27 | 4138 | 4475 | SqCL | AusARG (this project) |
| Pygopodidae_Aprasia_repens_WAMR153935 | 328 | 28 | 4255 | 4611 | SqCL | AusARG (this project) |
| Pygopodidae_Aprasia_repens_WAMR168645 | 357 | 32 | 4739 | 5128 | SqCL | Singhal et al. 2021 |
| Pygopodidae_Aprasia_repens_WAMR177945 | 314 | 28 | 4115 | 4457 | SqCL | AusARG (this project) |
| Pygopodidae_Aprasia_rostrata_153829 | 17 | 0 | 4549 | 4566 | UCE | Skipwith et al. 2019 |
| Pygopodidae_Aprasia_rostrata_WAMR130221 | 321 | 29 | 4468 | 4818 | SqCL | AusARG (this project) |
| Pygopodidae_Aprasia_rostrata_WAMR165984 | 337 | 30 | 4459 | 4826 | SqCL | AusARG (this project) |
| Pygopodidae_Aprasia_smithi_116919 | 17 | 0 | 4431 | 4448 | UCE | Skipwith et al. 2019 |
| Pygopodidae_Aprasia_smithi_WAMR116657 | 319 | 29 | 4081 | 4429 | SqCL | AusARG (this project) |
| Pygopodidae_Aprasia_smithi_WAMR116919 | 304 | 29 | 4212 | 4545 | SqCL | AusARG (this project) |
| Pygopodidae_Aprasia_striolata_156936 | 17 | 0 | 4439 | 4456 | UCE | Skipwith et al. 2019 |
| Pygopodidae_Aprasia_striolata_ABTC113417 | 329 | 28 | 4322 | 4679 | SqCL | AusARG (this project) |
| Pygopodidae_Aprasia_striolata_WAMR114483 | 295 | 29 | 4075 | 4399 | SqCL | AusARG (this project) |
| Pygopodidae_Aprasia_wicherina_WAMR121129 | 311 | 29 | 4335 | 4675 | SqCL | AusARG (this project) |
| Pygopodidae_Delma_australis_172541 | 15 | 0 | 4246 | 4261 | UCE | Skipwith et al. 2019 |
| Pygopodidae_Delma_australis_SAMAR39606 | 304 | 26 | 4152 | 4482 | SqCL | AusARG (this project) |
| Pygopodidae_Delma_australis_SAMAR50181 | 314 | 28 | 4238 | 4580 | SqCL | AusARG (this project) |
| Pygopodidae_Delma_australis_SAMAR60058 | 313 | 30 | 4150 | 4493 | SqCL | AusARG (this project) |
| Pygopodidae_Delma_australis_WAMR132470 | 167 | 14 | 1816 | 1997 | SqCL | AusARG (this project) |
| Pygopodidae_Delma_australis_WAMR175505 | 275 | 19 | 2571 | 2865 | SqCL | AusARG (this project) |
| Pygopodidae_Delma_australis_WAMR175657 | 167 | 11 | 1209 | 1387 | SqCL | AusARG (this project) |
| Pygopodidae_Delma_borea_ABTC057592 | 327 | 0 | 1 | 328 | AHE | Burbrink et al. 2021 |
| Pygopodidae_Delma_borea_CCM3160 | 17 | 0 | 4399 | 4416 | UCE | Skipwith et al. 2019 |
| Pygopodidae_Delma_borea_CCM3213 | 16 | 0 | 4252 | 4268 | UCE | Skipwith et al. 2019 |
| Pygopodidae_Delma_borea_MAGNTR22753 | 299 | 30 | 4200 | 4529 | SqCL | AusARG (this project) |
| Pygopodidae_Delma_borea_QMJ90130 | 257 | 25 | 3061 | 3343 | SqCL | AusARG (this project) |
| Pygopodidae_Delma_borea_SAMAR42019 | 286 | 29 | 3979 | 4294 | SqCL | AusARG (this project) |
| Pygopodidae_Delma_borea_WAMR165969 | 206 | 18 | 1789 | 2013 | SqCL | AusARG (this project) |
| Pygopodidae_Delma_butleri_R35031 | 16 | 0 | 4632 | 4648 | UCE | Skipwith et al. 2019 |
| Pygopodidae_Delma_butleri_SAMAR35033 | 266 | 28 | 3965 | 4259 | SqCL | AusARG (this project) |
| Pygopodidae_Delma_butleri_UMMZ21111 | 295 | 25 | 4237 | 4557 | SqCL | AusARG (this project) |
| Pygopodidae_Delma_butleri_UMMZ21172 | 300 | 28 | 3982 | 4310 | SqCL | AusARG (this project) |
| Pygopodidae_Delma_butleri_UMMZ244261 | 320 | 29 | 4785 | 5134 | SqCL | Title & Singhal et al. 2024 |
| Pygopodidae_Delma_butleri_WAMR120819 | 334 | 26 | 4640 | 5000 | SqCL | AusARG (this project) |
| Pygopodidae_Delma_concinna_ABTC113062 | 229 | 24 | 2692 | 2945 | SqCL | AusARG (this project) |
| Pygopodidae_Delma_concinna_WAMR141175 | 244 | 26 | 3640 | 3910 | SqCL | AusARG (this project) |
| Pygopodidae_Delma_concinnna_ABTC113063 | 247 | 26 | 3345 | 3618 | SqCL | AusARG (this project) |
| Pygopodidae_Delma_desmosa_166043 | 16 | 0 | 4419 | 4435 | UCE | Skipwith et al. 2019 |
| Pygopodidae_Delma_desmosa_WAMR134414 | 305 | 27 | 4105 | 4437 | SqCL | AusARG (this project) |
| Pygopodidae_Delma_desmosa_WAMR163958 | 274 | 27 | 3827 | 4128 | SqCL | AusARG (this project) |
| Pygopodidae_Delma_desmosa_WAMR175439 | 342 | 30 | 4698 | 5070 | SqCL | AusARG (this project) |
| Pygopodidae_Delma_elegans_RL93 | 18 | 0 | 4284 | 4302 | UCE | Skipwith et al. 2019 |
| Pygopodidae_Delma_elegans_WAMR163210 | 299 | 26 | 3763 | 4088 | SqCL | AusARG (this project) |
| Pygopodidae_Delma_fraseri_169904 | 17 | 0 | 4585 | 4602 | UCE | Skipwith et al. 2019 |
| Pygopodidae_Delma_fraseri_WAMR175607 | 298 | 25 | 4343 | 4666 | SqCL | AusARG (this project) |

Table S1: Table of molecular samples. Fields indicate the number of loci recovered per sample, as well as the project under which the data were generated. *(continued)*

| lineage | AHE | gene | uce | total_loci | Targets | Source |
|---|---|---|---|---|---|---|
| Pygopodidae__Delma__grayii__154064 | 17 | 0 | 4613 | 4630 | UCE | Skipwith et al. 2019 |
| Pygopodidae__Delma__grayii__WAMR131871 | 282 | 29 | 4061 | 4372 | SqCL | AusARG (this project) |
| Pygopodidae__Delma__grayii__WAMR146399 | 114 | 8 | 1033 | 1155 | SqCL | AusARG (this project) |
| Pygopodidae__Delma__haroldi__172326 | 16 | 0 | 4597 | 4613 | UCE | Skipwith et al. 2019 |
| Pygopodidae__Delma__haroldi__WAMR154780 | 306 | 26 | 4224 | 4556 | SqCL | AusARG (this project) |
| Pygopodidae__Delma__haroldi__WAMR156703 | 282 | 26 | 3698 | 4006 | SqCL | AusARG (this project) |
| Pygopodidae__Delma__hebesa__WAMR175526 | 295 | 25 | 4101 | 4421 | SqCL | AusARG (this project) |
| Pygopodidae__Delma__hebesa__WAMR175581 | 342 | 31 | 4676 | 5049 | SqCL | AusARG (this project) |
| Pygopodidae__Delma__impar__ABTC17683 | 219 | 21 | 2224 | 2464 | SqCL | AusARG (this project) |
| Pygopodidae__Delma__impar__ABTC68788 | 278 | 24 | 3413 | 3715 | SqCL | AusARG (this project) |
| Pygopodidae__Delma__inornata__ABTC144849 | 291 | 27 | 3554 | 3872 | SqCL | AusARG (this project) |
| Pygopodidae__Delma__inornata__SAMAR36585 | 312 | 28 | 4386 | 4726 | SqCL | AusARG (this project) |
| Pygopodidae__Delma__inornata__SAMAR44457 | 315 | 28 | 4288 | 4631 | SqCL | AusARG (this project) |
| Pygopodidae__Delma__labialis__QMJ89155 | 287 | 27 | 3944 | 4258 | SqCL | AusARG (this project) |
| Pygopodidae__Delma__labialis__QMJ89591 | 313 | 28 | 4253 | 4594 | SqCL | AusARG (this project) |
| Pygopodidae__Delma__labialis__SMZ__1489 | 312 | 24 | 4121 | 4457 | SqCL | AusARG (this project) |
| Pygopodidae__Delma__mitella__ABTC58998 | 267 | 25 | 3670 | 3962 | SqCL | AusARG (this project) |
| Pygopodidae__Delma__molleri__ABTC131082 | 319 | 28 | 4351 | 4698 | SqCL | AusARG (this project) |
| Pygopodidae__Delma__molleri__ABTC136854 | 314 | 27 | 4397 | 4738 | SqCL | AusARG (this project) |
| Pygopodidae__Delma__nasuta__AMSR.118632 | 317 | 27 | 4422 | 4766 | SqCL | AusARG (this project) |
| Pygopodidae__Delma__nasuta__CCM3360 | 16 | 0 | 4281 | 4297 | UCE | Skipwith et al. 2019 |
| Pygopodidae__Delma__nasuta__MAGNTR35901 | 335 | 31 | 4687 | 5053 | SqCL | AusARG (this project) |
| Pygopodidae__Delma__nasuta__WAMR133433 | 284 | 27 | 4157 | 4468 | SqCL | AusARG (this project) |
| Pygopodidae__Delma__pax__172570 | 17 | 0 | 4543 | 4560 | UCE | Skipwith et al. 2019 |
| Pygopodidae__Delma__pax__WAMR175347 | 287 | 23 | 3931 | 4241 | SqCL | AusARG (this project) |
| Pygopodidae__Delma__petersoni__166727 | 18 | 0 | 4534 | 4552 | UCE | Skipwith et al. 2019 |
| Pygopodidae__Delma__petersoni__WAMR165873 | 309 | 28 | 4288 | 4625 | SqCL | AusARG (this project) |
| Pygopodidae__Delma__petersoni__WAMR165874 | 282 | 27 | 3858 | 4167 | SqCL | AusARG (this project) |
| Pygopodidae__Delma__plebeia__QMJ86597 | 343 | 29 | 0 | 372 | SqCL | AusARG (this project) |
| Pygopodidae__Delma__tealei__153819 | 17 | 0 | 4595 | 4612 | UCE | Skipwith et al. 2019 |
| Pygopodidae__Delma__tealei__WAMR153811 | 309 | 30 | 4369 | 4708 | SqCL | AusARG (this project) |
| Pygopodidae__Delma__tealei__WAMR153819 | 269 | 26 | 3658 | 3953 | SqCL | AusARG (this project) |
| Pygopodidae__Delma__tincta__ABTC60820 | 338 | 31 | 4658 | 5027 | SqCL | AusARG (this project) |
| Pygopodidae__Delma__tincta__MAGNTR22445 | 273 | 27 | 3715 | 4015 | SqCL | AusARG (this project) |
| Pygopodidae__Delma__tincta__NTMR37588 | 252 | 27 | 3686 | 3965 | SqCL | AusARG (this project) |
| Pygopodidae__Delma__tincta__QMJ80136 | 341 | 31 | 4691 | 5063 | SqCL | AusARG (this project) |
| Pygopodidae__Delma__tincta__QMJ84129 | 205 | 27 | 3011 | 3243 | SqCL | AusARG (this project) |
| Pygopodidae__Delma__tincta__QMJ93148 | 240 | 27 | 3504 | 3771 | SqCL | AusARG (this project) |
| Pygopodidae__Delma__tincta__QMJ94263 | 338 | 29 | 4778 | 5145 | SqCL | AusARG (this project) |
| Pygopodidae__Delma__tincta__SAMAR30971 | 339 | 29 | 4691 | 5059 | SqCL | AusARG (this project) |
| Pygopodidae__Delma__tincta__SAMAR40238 | 342 | 30 | 4688 | 5060 | SqCL | AusARG (this project) |
| Pygopodidae__Delma__tincta__SAMAR55294 | 337 | 30 | 4682 | 5049 | SqCL | AusARG (this project) |
| Pygopodidae__Delma__tincta__SAMAR60966 | 303 | 26 | 4263 | 4592 | SqCL | AusARG (this project) |
| Pygopodidae__Delma__tincta__UMMZ242601 | 349 | 30 | 4727 | 5106 | SqCL | Title & Singhal et al. 2024 |
| Pygopodidae__Delma__tincta__UMMZ242601.2 | 347 | 29 | 4692 | 5068 | SqCL | AusARG (this project) |
| Pygopodidae__Delma__tincta__WAMR114391 | 341 | 31 | 4746 | 5118 | SqCL | AusARG (this project) |
| Pygopodidae__Delma__tincta__WAMR157124 | 326 | 28 | 4618 | 4972 | SqCL | AusARG (this project) |
| Pygopodidae__Delma__tincta__WAMR168506 | 330 | 29 | 4723 | 5082 | SqCL | AusARG (this project) |
| Pygopodidae__Delma__tincta__WAMR168506.1 | 304 | 23 | 4368 | 4695 | SqCL | AusARG (this project) |
| Pygopodidae__Delma__torquata__QMJ83187 | 342 | 28 | 4495 | 4865 | SqCL | AusARG (this project) |
| Pygopodidae__Delma__torquata__QMJ93842 | 345 | 27 | 4633 | 5005 | SqCL | AusARG (this project) |
| Pygopodidae__Delma__vescolineata__ABTC11106 | 158 | 23 | 1853 | 2034 | SqCL | AusARG (this project) |
| Pygopodidae__Lialis__burtonis__ABTC29938 | 235 | 15 | 1606 | 1856 | SqCL | AusARG (this project) |
| Pygopodidae__Lialis__burtonis__ABTC30417 | 302 | 27 | 4175 | 4504 | SqCL | AusARG (this project) |
| Pygopodidae__Lialis__burtonis__ABTC3721 | 208 | 14 | 1828 | 2050 | SqCL | AusARG (this project) |
| Pygopodidae__Lialis__burtonis__ABTC52145 | 323 | 26 | 4275 | 4624 | SqCL | AusARG (this project) |
| Pygopodidae__Lialis__burtonis__ABTC52146 | 293 | 23 | 4102 | 4418 | SqCL | AusARG (this project) |
| Pygopodidae__Lialis__burtonis__ABTC58538 | 270 | 28 | 3918 | 4216 | SqCL | AusARG (this project) |
| Pygopodidae__Lialis__burtonis__ABTC70465 | 323 | 28 | 4374 | 4725 | SqCL | AusARG (this project) |
| Pygopodidae__Lialis__burtonis__BPBM35951 | 338 | 0 | 1 | 339 | AHE | Burbrink et al. 2021 |
| Pygopodidae__Lialis__burtonis__CCM3273 | 18 | 0 | 4219 | 4237 | UCE | Skipwith et al. 2019 |
| Pygopodidae__Lialis__burtonis__RJE110407 | 336 | 30 | 4584 | 4950 | SqCL | AusARG (this project) |
| Pygopodidae__Lialis__burtonis__RJE__110407 | 309 | 25 | 4203 | 4537 | SqCL | AusARG (this project) |
| Pygopodidae__Lialis__burtonis__UMMZ244266 | 315 | 29 | 4790 | 5134 | SqCL | Title & Singhal et al. 2024 |
| Pygopodidae__Lialis__jicari__LSUMZ10483 | 17 | 0 | 4272 | 4289 | UCE | Skipwith et al. 2019 |
| Pygopodidae__Ophidiocephalus__taeniatus__SAMAR28366 | 347 | 32 | 4761 | 5140 | SqCL | AusARG (this project) |
| Pygopodidae__Ophidiocephalus__taeniatus__SAMAR28366.2 | 328 | 29 | 4552 | 4909 | SqCL | AusARG (this project) |
| Pygopodidae__Paradelma__orientalis__PMO230 | 348 | 29 | 4727 | 5104 | SqCL | AusARG (this project) |
| Pygopodidae__Paradelma__orientalis__PMO__230 | 320 | 25 | 4414 | 4759 | SqCL | AusARG (this project) |
| Pygopodidae__Paradelma__orientalis__QMJ92675 | 17 | 0 | 4628 | 4645 | UCE | Skipwith et al. 2019 |
| Pygopodidae__Pletholax__edelensis__WAMR120973 | 304 | 29 | 4101 | 4434 | SqCL | AusARG (this project) |
| Pygopodidae__Pletholax__edelensis__WAMR156967 | 300 | 29 | 4091 | 4420 | SqCL | AusARG (this project) |

Table S1: Table of molecular samples. Fields indicate the number of loci recovered per sample, as well as the project under which the data were generated. *(continued)*

| lineage | AHE | gene | uce | total_loci | Targets | Source |
|---|---|---|---|---|---|---|
| Pygopodidae__Pletholax__gracilis__106171 | 17 | 0 | 4587 | 4604 | UCE | Skipwith et al. 2019 |
| Pygopodidae__Pletholax__gracilis__WAMR119115 | 288 | 28 | 3973 | 4289 | SqCL | AusARG (this project) |
| Pygopodidae__Pletholax__gracilis__WAMR154023 | 344 | 30 | 4719 | 5093 | SqCL | AusARG (this project) |
| Pygopodidae__Pletholax__gracilis__WAMR154023 2 | 304 | 27 | 4237 | 4568 | SqCL | AusARG (this project) |
| Pygopodidae__Pygopus__lepidopodus__ABTC11338 | 317 | 25 | 4251 | 4593 | SqCL | AusARG (this project) |
| Pygopodidae__Pygopus__lepidopodus__ABTC34649 | 321 | 28 | 4294 | 4643 | SqCL | AusARG (this project) |
| Pygopodidae__Pygopus__lepidopodus__RL87 | 16 | 0 | 4358 | 4374 | UCE | Skipwith et al. 2019 |
| Pygopodidae__Pygopus__nigriceps__CUMV14267 | 348 | 30 | 4814 | 5192 | SqCL | Title & Singhal et al. 2024 |
| Pygopodidae__Pygopus__nigriceps__R35761 | 14 | 0 | 4283 | 4297 | UCE | Skipwith et al. 2019 |
| Pygopodidae__Pygopus__nigriceps__RL204 | 17 | 0 | 4317 | 4334 | UCE | Skipwith et al. 2019 |
| Pygopodidae__Pygopus__nigriceps__WAMR136743 | 342 | 31 | 4679 | 5052 | SqCL | AusARG (this project) |
| Pygopodidae__Pygopus__nigriceps__WAMR136743 2 | 303 | 29 | 4306 | 4638 | SqCL | AusARG (this project) |
| Pygopodidae__Pygopus__robertsi__ABTC28230 | 306 | 24 | 4297 | 4627 | SqCL | AusARG (this project) |
| Pygopodidae__Pygopus__schraderi__ABTC118373 | 311 | 26 | 4365 | 4702 | SqCL | AusARG (this project) |
| Pygopodidae__Pygopus__schraderi__ABTC127627 | 95 | 6 | 606 | 707 | SqCL | AusARG (this project) |
| Pygopodidae__Pygopus__schraderi__ABTC72730 | 93 | 2 | 947 | 1042 | SqCL | AusARG (this project) |
| Pygopodidae__Pygopus__schraderi__QMJ77344 | 343 | 29 | 4632 | 5004 | SqCL | AusARG (this project) |
| Pygopodidae__Pygopus__schraderi__S90 | 17 | 0 | 4439 | 4456 | UCE | Skipwith et al. 2019 |
| Pygopodidae__Pygopus__steelescotti__ABTC29348 | 258 | 23 | 2967 | 3248 | SqCL | AusARG (this project) |
| Pygopodidae__Pygopus__steelescotti__ABTC30411 | 333 | 28 | 4593 | 4954 | SqCL | AusARG (this project) |
| Pygopodidae__Pygopus__steelescotti__R35022 | 17 | 0 | 4562 | 4579 | UCE | Skipwith et al. 2019 |
| Pygopodidae__Pygopus__steelescotti__WAMR164760 | 134 | 9 | 1575 | 1718 | SqCL | AusARG (this project) |

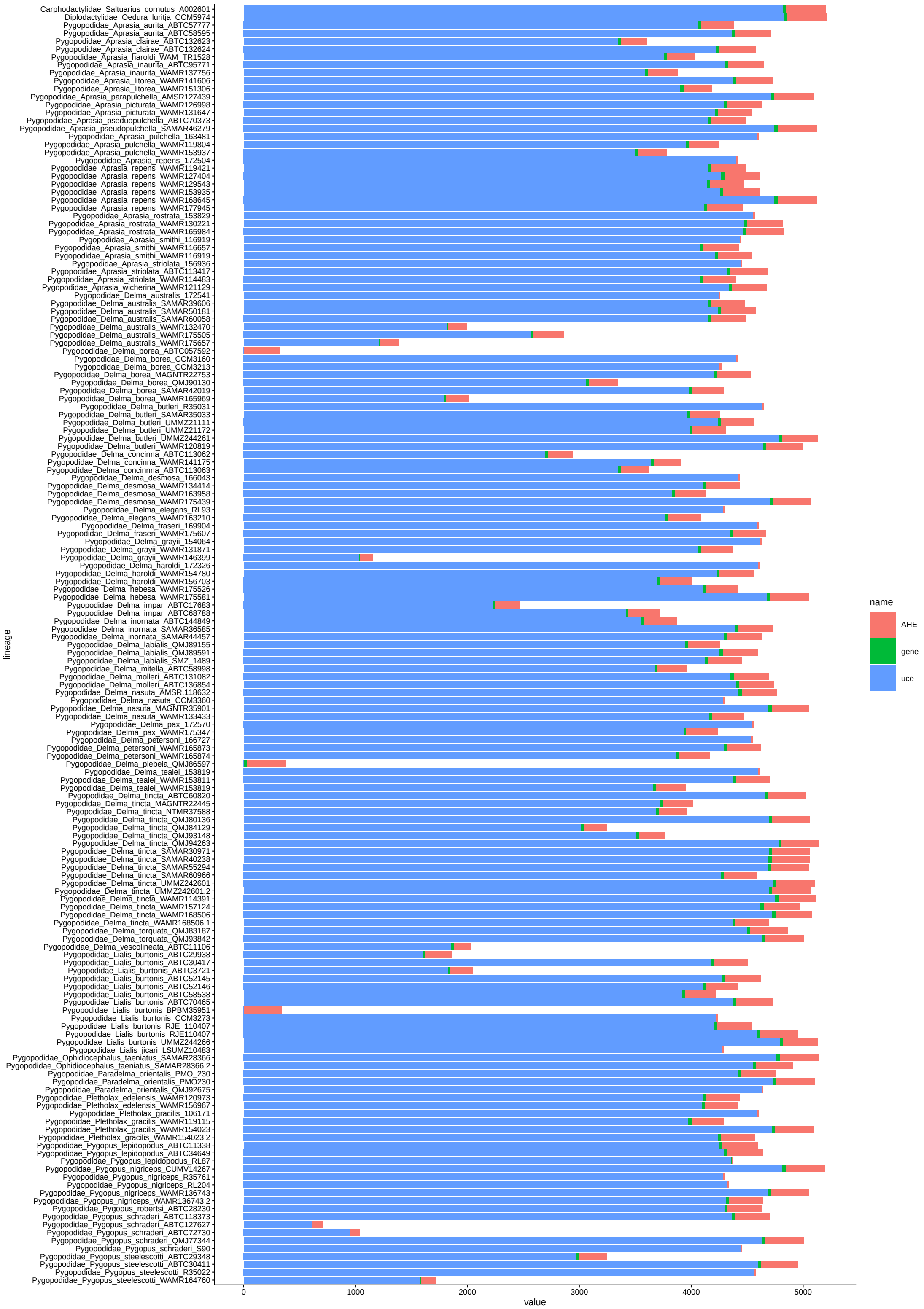

**Figure S1.** Locus type and number summarized by sample for newly sequenced samples. Dotted line indicates 50% of targeted sequences recovered. Samples in grey text at left were not included in final analyses.

##### Phylogenetic Analyses

We estimated individual genealogies for our sequence-capture data (n=5441) under maximum-likelihood in IQ-TREE2 (Minh et al. 2020) allowing the program to assign the best fitting substitution model using ModelFinder (Kalyaanamoorthy et al. 2017), then perform 1,000 ultrafast bootstraps (Minh et al. 2013). We then estimated the species tree using the coalescent-consistent methods wASTRAL-hybrid and ASTRAL-IV (Zhang & Mirarab 2022), using IQ-TREE2 gene trees as input. We used two ASTRAL methods to take advantage of the way wASTRAL-hybrid weights branch lengths and support values in estimating the species tree, and how ASTRAL-IV allows multiple individuals per species and estimates both internal and terminal branch lengths in substitutions-per-site (as opposed to coalescent units for internal branch lengths and static terminal branch lengths in wASTRAL). To quantify topological signal from individual gene trees we calculated gene concordance factors using IQ-TREE2 and focused on intergeneric relationships using a single representative for each genus.

To incorporate our dated pygopodid tree into the squamate framework of Title et al. (2024) we started by selecting their best chosen fully sampled tree (`best_ultrametric_fulltree_ddBD_`[illegible] We dropped all pygopodid tips, bar one, then grafted our tree onto that tip at the height of the estimated pygopodid crown using `phytools`. This tree is included as Supplement/Trees/Title_Tree.tre.

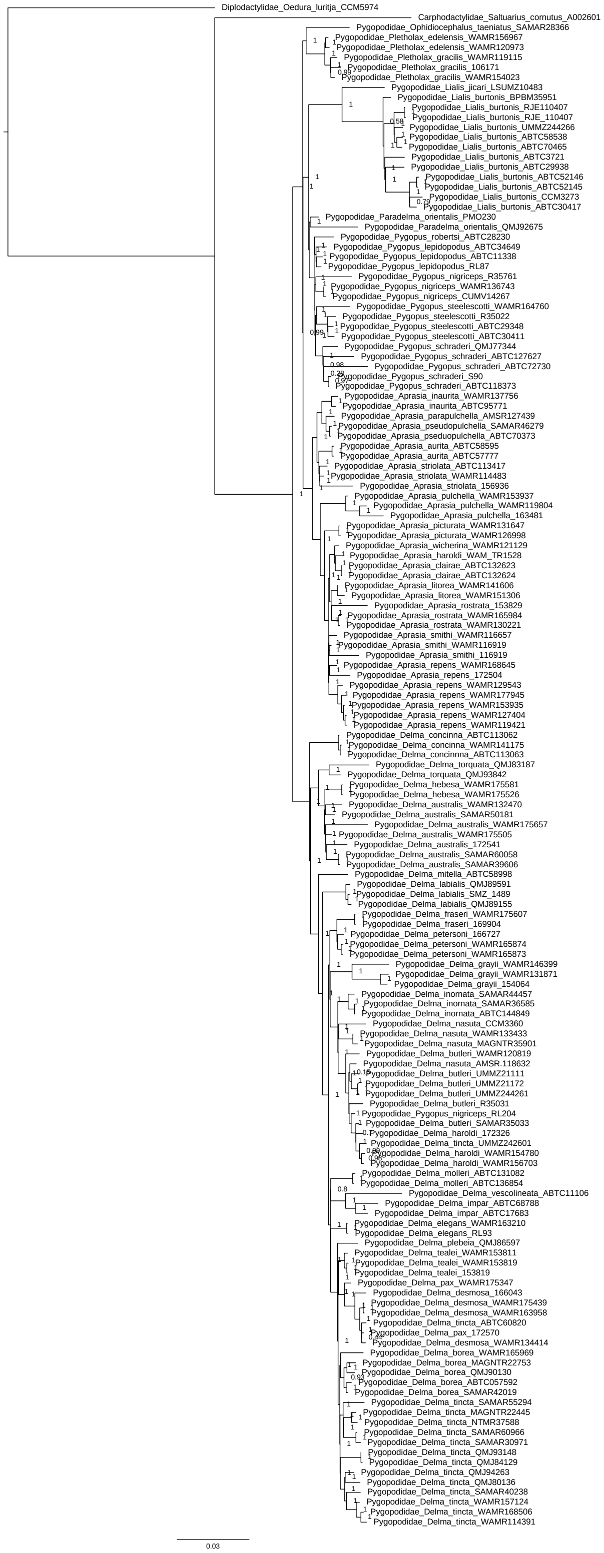

**Figure S2.** Fully sampled sequence capture tree, estimated by ASTRAL4 using IQTREE-2 genetree inputs. Branch support values are indicated at nodes.

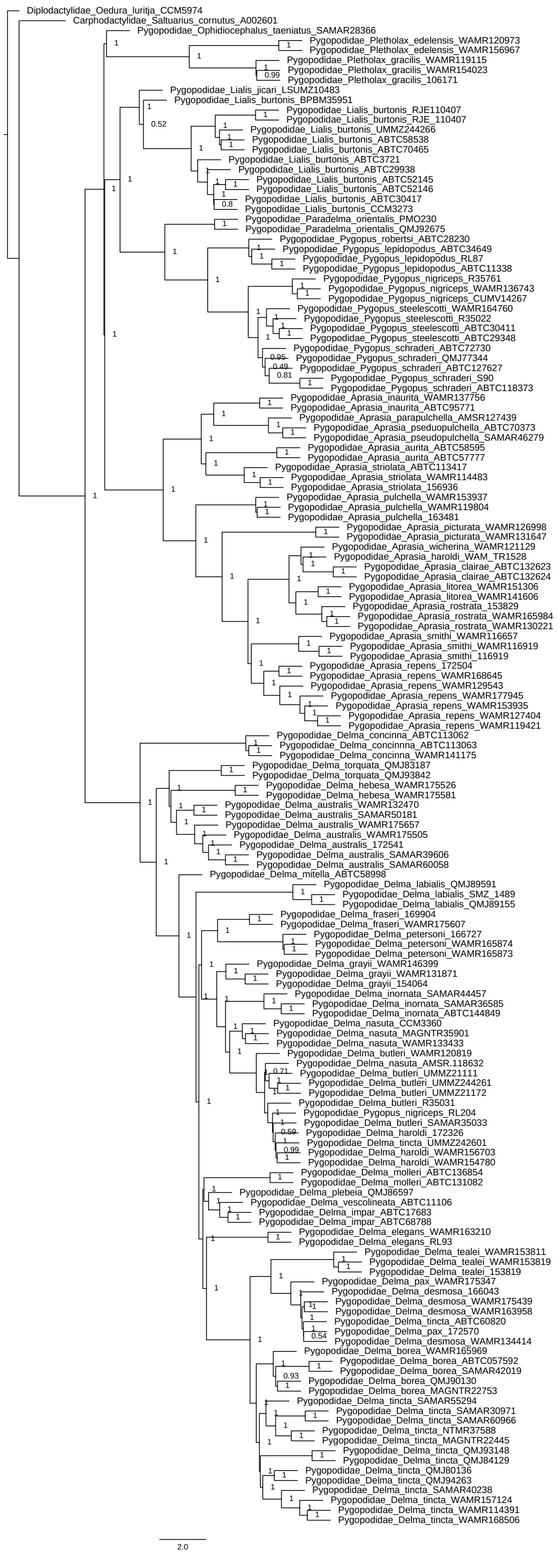

**Figure S3.** Fully sampled sequence capture tree, estimated by weighted ASTRAL hybrid using IQTREE-2 genetree inputs. Branch support values are indicated at nodes.

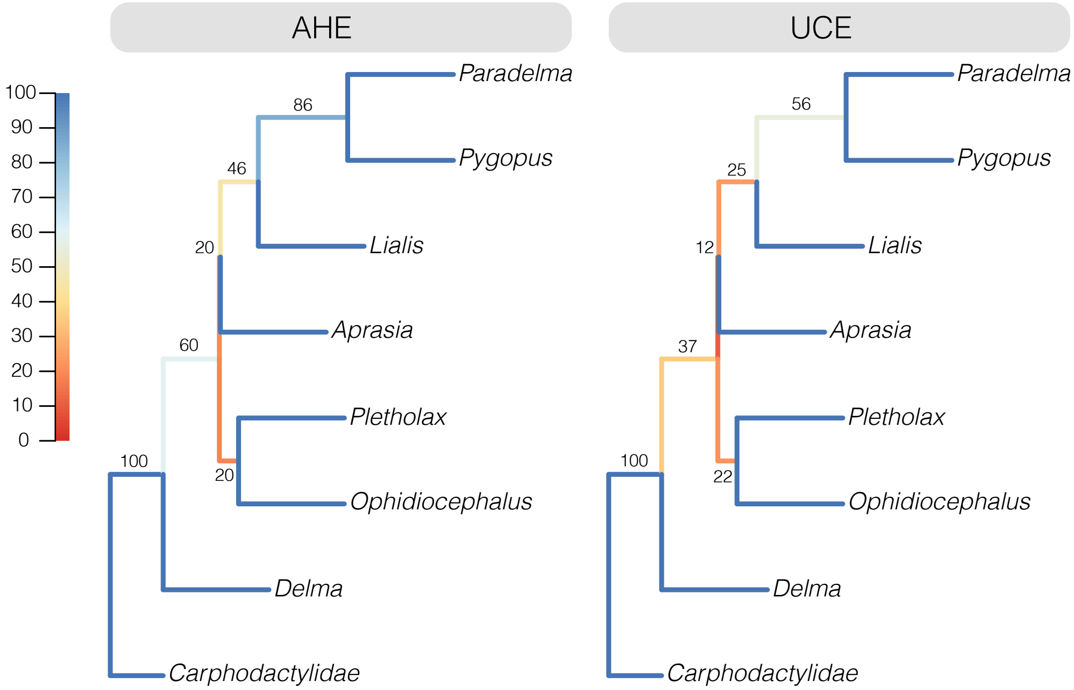

**Figure S4.** Low gene concordance factors (gCF) among some genera indicate moderate levels of incomplete lineage sorting. gCF scores are typically higher across the AHE dataset relative to the UCE dataset.

#### Divergence Dating

To estimate divergence times among taxa we applied a series of fossil and secondary calibrations in MCMCTree (Rannala & Yang 2007) as outlined in the Supplement and Table S2. We started by trimming our ASTRAL tree down to a single representative for each ingroup species or subspecies for input. We reduced our molecular data to exonic markers in the AHE loci, then estimated raw genetic distances from alignments to use as a proxy for evolutionary rate. We removed the fastest 5% and slowest 5% of loci to avoid issues with extreme rate heterogeneity, divided the remaining loci into four rate partitions, and then removed third codon positions from all loci using AMAS (Borowiec 2016).

For each partition we ran MCMCTree with usedata = 3 to get the approximate likelihoods and branch lengths using baseml (dos Reis & Yang 2011), then concatenated the out.BV files together. We then ran four replicate MCMCTree analyses on the gradient and Hessian (in.BV file; usedata = 2), each for 20k burnin generations before collecting 20k samples at a sampling frequency of 100 generations (2,020,000 total generations). We compared mcmc files for stationarity and convergence (ESS of all parameters > 200), combined them using logCombiner, and used this combined mcmc file to summarize divergence times on our tree (print = -1 in .ctl file). To validate our priors we ran an additional analysis (usedata = 0) to run explicitly from the prior calibrations and determine our effective priors for comparison against our posterior age estimates. We then plotted the applied priors against effective priors (priors as a result of multiple interacting priors from usedata = 0) and posterior estimates to ensure appropriate behavior of the MCMCTree analyses.

**Table S2. Fossil Calibrations**

| Cal. | Fossil Information | Calibration | Split/Position | Source |
|---|---|---|---|---|
| A | Uniform—Sophineta | 'B(2.38,2.55)' | Lepidosauria (Sphenodon + Squamata) | see below |
| B | Uniform—Secondary | 'B(0.50,0.80)' | Diplodactylidae + (Carphodactylidae + Pygopodidae) | Burbrink et al. 2020; Title et al. 2024 |

*Note.* The root divergence between lepidosaurs (*Sphenodon* + squamates) is based on the fossil taxa *Sophineta cracoviensis* (Evans & Bialynicka, 2009), *Megachirella wachtleri* (Renesto & Posenato, 2003), and the Vellberg Jaw (Jones et al., 2013). These taxa represent stem squamates and rhynchocephalians, and so provide a soft lower bound on the crown divergence of Lepidosauria, with a soft upper bound provided by *Protosaurus speneri* (von Meyer, 1832).

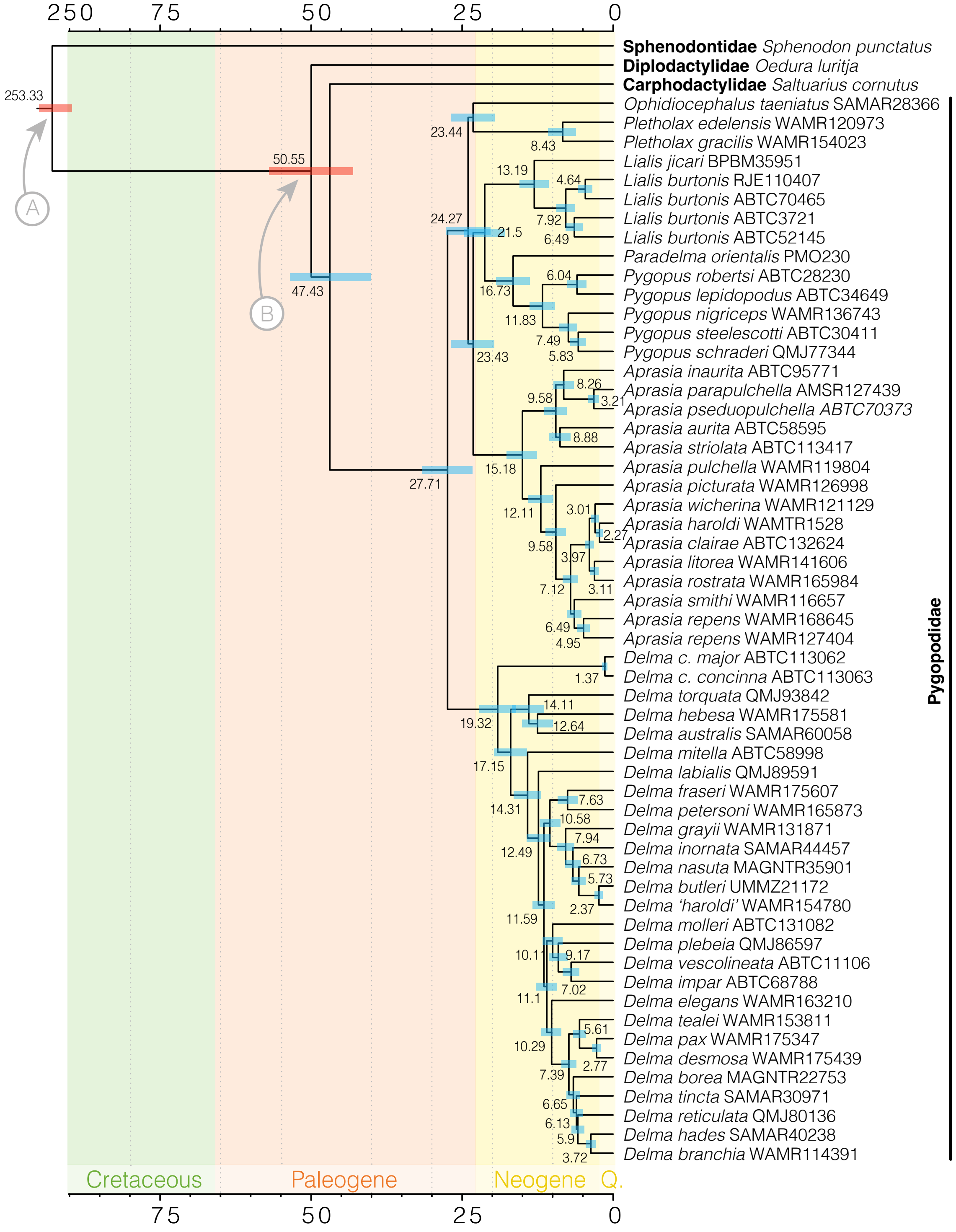

**Figure S5.** Fully sampled time-tree estimated using MCMCTree. Divergence estimates are indicated at each node and accompanied by colored bars to show 95% confidence intervals. Red node bars and grey arrows highlight calibrated nodes outlined in Table S2.

### Morphological Sampling

#### Database Assembly

We collected two morphological traits traditionally measured in squamate research, Snout-Vent Length (SVL) and Tail Length (TL). When combined, these traits are referred to as Total Length (Total). SVL is measured from the tip of the snout to the cloaca. In small species this is typically done by straightening the body and using a pair of calipers. In larger species and in snakes, where measuring via calipers is not possible, a non-elastic and dampened piece of string is typically used to capture the length of the specimen and then it is compared against a measuring tape, ruler, or yardstick. TL is measured from the cloaca to the tip of the tail. Because many species of lizards have the ability to autotomize, or drop their tails, only original (non-regenerated) tails are measured. The process of measuring is typically the same as for SVL: by calipers in species with short tails and longer tails are measured by string. The morphological database of these traits is available in `Data/Head_Body_Tail.csv`. To understand the co-evolution of these two traits, we generated a composite measure, the ratio of tail-to-body length (TL/SVL).

Raw morphological measurements were collected from museum specimens by all three authors as part of the Australian Amphibian and Reptile Genomics initiative and an Australian Research Council grant to JSK (DP210100820), provided as unpublished data, and collated from published sources. Information about the data collector (where known) and data source are provided in the data file. A significant number of records come from published datasets, including: Anelli et al. 2024 (1,420 records), Bergmann & Morinaga 2019 (231), Brandley et al. 2008 (242), Brennan et al. 2026 (460), Grundler & Rabosky 2014 (300), Jennings 2002 (747 for pygopods), Norris et al. 2021 (1,189), Meiri 2024 (1,045), Sheehy III et al. 2014 (225), Wang et al. 2026 (322). Additionally, a large number represent previously unpublished efforts first presented here: AusARG/ARC DP Keogh et al. (1,178), Chapple Group (4,207). In combining datasets all records were cleaned manually by a series of filtering steps. These included (1) flagging and inspecting records of snakes with tail lengths that exceeded 75% of their snout-vent length, (2) visualizing trends in TL/SVL-to-Total length at a family level to identify outliers (Fig.S6), and (3) flagging and inspecting any records that fell outside two standard deviations of the mean of the family they belong to for TL/SVL. Flagged records that could not be independently verified were removed from the database.

To investigate trends in tail evolution across major squamate groups and families we fit linear models to each group/family with >3 species represented in the dataset (`Scripts/04_LinearModels.R`). Visualizations are presented in Figure S6, and details are in Table S3 and S4 below. Note, linear models are likely an inappropriate fit to some groups with highly varied ecologies (e.g. Lacertoidea, which contains long-tailed lacertids and short tailed amphisbaenians).

Table S3: Fit of linear model to each squamate group. Ordered by ascending slope.

| group | r2 | intercept | slope | std_err | p_value | n_spp |
|---|---|---|---|---|---|---|
| Lacertoidea | -0.0070542 | 1.4303403 | -0.0000677 | 0.0009282 | 0.9419675 | 143 |
| Snakes | 0.0624180 | 0.1643885 | 0.0000724 | 0.0000066 | 0.0000000 | 1801 |
| Anguimorpha | 0.0931421 | 1.2300015 | 0.0004636 | 0.0001630 | 0.0058837 | 70 |
| Dibamia | -0.0277336 | 0.0767758 | 0.0005438 | 0.0006365 | 0.4150009 | 11 |
| Xantusiia | -0.4820665 | 1.3138784 | 0.0010879 | 0.0018403 | 0.6601137 | 3 |
| Scincidae | 0.0389786 | 1.0141596 | 0.0014021 | 0.0002715 | 0.0000003 | 634 |
| Iguania | 0.2638006 | 1.4008651 | 0.0022158 | 0.0002800 | 0.0000000 | 173 |
| Gekkota | 0.4357593 | 0.3454848 | 0.0041568 | 0.0002796 | 0.0000000 | 286 |
| Cordyliformes | 0.3906274 | 0.9610412 | 0.0043919 | 0.0009469 | 0.0000604 | 33 |

Table S4: Fit of linear model to each squamate family. Ordered by ascending slope.

| group | family | r2 | intercept | slope | std_err | p_value | n_spp |
|---|---|---|---|---|---|---|---|
| Gekkota | Eublepharidae | -0.7910092 | 0.8244558 | -0.0016133 | 0.0047228 | 0.7904440 | 3 |
| Snakes | Tropidophiidae | 0.3464356 | 0.2605209 | -0.0003371 | 0.0001553 | 0.0730249 | 8 |
| Snakes | Leptotyphlopidae | -0.0000258 | 0.1456855 | -0.0002558 | 0.0002559 | 0.3257662 | 31 |
| Lacertoidea | Amphisbaenidae | 0.3138029 | 0.1653799 | -0.0002047 | 0.0000777 | 0.0217603 | 14 |
| Lacertoidea | Amphisbaenia | 0.0283570 | 0.1538946 | -0.0001918 | 0.0001669 | 0.2771540 | 12 |
| Snakes | Atractaspididae | -0.0305629 | 0.2244193 | -0.0001189 | 0.0001688 | 0.4914567 | 18 |
| Snakes | Uropeltidae | 0.0086726 | 0.0884444 | -0.0001095 | 0.0001006 | 0.2895381 | 22 |
| Snakes | Anomalepididae | -0.1542244 | 0.0985654 | -0.0000822 | 0.0003231 | 0.8077401 | 8 |
| Snakes | Pseudoxyrhophiidae | -0.1081289 | 0.3628010 | -0.0000814 | 0.0001265 | 0.5480433 | 7 |
| Snakes | Xenopeltidae | 0.5695538 | 0.1411505 | -0.0000647 | 0.0000339 | 0.3071160 | 3 |
| Snakes | Calamariidae | -0.4798013 | 0.1126762 | -0.0000450 | 0.0002721 | 0.8839578 | 4 |
| Snakes | Psammophiidae | -0.0264009 | 0.3940890 | -0.0000279 | 0.0000716 | 0.6999862 | 34 |
| Snakes | Cylindrophiidae | -0.1986979 | 0.0252586 | -0.0000014 | 0.0000186 | 0.9441090 | 7 |
| Snakes | Pythonidae | -0.1248159 | 0.1488999 | -0.0000009 | 0.0000240 | 0.9720193 | 10 |
| Snakes | Natricidae | -0.0250000 | 0.3069352 | 0.0000000 | 0.0001097 | 0.9997116 | 42 |
| Snakes | Typhlopidae | -0.0101231 | 0.0292880 | 0.0000074 | 0.0000265 | 0.7806458 | 93 |
| Snakes | Viperids | -0.0968108 | 0.1177345 | 0.0000107 | 0.0000626 | 0.8680013 | 12 |
| Snakes | Henophidiae | 0.1409248 | 0.0780438 | 0.0000259 | 0.0000064 | 0.0000970 | 96 |
| Snakes | Elapidae | 0.0356755 | 0.1290565 | 0.0000270 | 0.0000081 | 0.0010224 | 272 |
| Snakes | Boidae | 0.0887334 | 0.1121316 | 0.0000317 | 0.0000182 | 0.0963415 | 22 |
| Snakes | Xenodermidae | -0.0238420 | 0.2490898 | 0.0000620 | 0.0001051 | 0.5601693 | 29 |
| Snakes | Gerrhopilidae | -0.0074560 | 0.0155381 | 0.0000742 | 0.0000765 | 0.3643862 | 9 |
| Anguimorpha | Varanidae | -0.0249439 | 1.4784005 | 0.0000798 | 0.0001798 | 0.6602348 | 34 |
| Snakes | Viperidae | 0.0300489 | 0.1267658 | 0.0000835 | 0.0000375 | 0.0276184 | 129 |
| Snakes | Ungaliophiidae | -0.6136590 | 0.0966043 | 0.0000875 | 0.0001788 | 0.7103030 | 3 |
| Snakes | Lamprophiidae | 0.1437560 | 0.1194148 | 0.0001207 | 0.0000334 | 0.0005526 | 73 |
| Snakes | Colubridae | 0.1292015 | 0.1894466 | 0.0001266 | 0.0000116 | 0.0000000 | 791 |
| Lacertoidea | Teiidae | -0.1382043 | 2.0713186 | 0.0001785 | 0.0002848 | 0.5647843 | 6 |
| Lacertoidea | Blanidae | -0.4225835 | 0.0875672 | 0.0002023 | 0.0006132 | 0.7728195 | 4 |
| Snakes | Homalopsids | 0.4929900 | 0.0333275 | 0.0002506 | 0.0001266 | 0.1863703 | 4 |
| Snakes | Pareidae | 0.1115371 | 0.0560334 | 0.0003769 | 0.0001623 | 0.0263283 | 36 |
| Snakes | Pseudoxenodontidae | 0.6779614 | -0.0253038 | 0.0004009 | 0.0001482 | 0.1138242 | 4 |
| Snakes | Homalopsidae | 0.4594344 | -0.0432706 | 0.0004899 | 0.0001411 | 0.0046211 | 14 |
| Dibamia | Dibamidae | -0.0277336 | 0.0767758 | 0.0005438 | 0.0006365 | 0.4150009 | 11 |
| Gekkota | Gekkonidae | 0.0235129 | 0.9346571 | 0.0006014 | 0.0003545 | 0.0938275 | 79 |
| Snakes | Prosymnidae | 0.1832169 | -0.0762897 | 0.0006253 | 0.0004539 | 0.2621601 | 5 |
| Xantusiia | Xantusiidae | -0.4820665 | 1.3138784 | 0.0010879 | 0.0018403 | 0.6601137 | 3 |
| Iguania | Iguanidae | -0.3050343 | 1.8288713 | 0.0011648 | 0.0015962 | 0.5986679 | 3 |
| Anguimorpha | Anguidae | 0.2882876 | 1.0753522 | 0.0012584 | 0.0003773 | 0.0027606 | 26 |
| Scincidae | Scincidae | 0.0389786 | 1.0141596 | 0.0014021 | 0.0002715 | 0.0000003 | 634 |
| Anguimorpha | Diploglossidae | 0.3644519 | 0.8104439 | 0.0015997 | 0.0008815 | 0.1671737 | 5 |
| Gekkota | Diplodactylidae | 0.1240909 | 0.4787039 | 0.0019144 | 0.0004566 | 0.0000542 | 118 |
| Iguania | Agamidae | 0.2416421 | 1.5657358 | 0.0019360 | 0.0003030 | 0.0000000 | 126 |
| Cordyliformes | Gerrhosauridae | -0.0148219 | 1.6667056 | 0.0019719 | 0.0022315 | 0.3917834 | 16 |
| Iguania | Chamaeleonidae | 0.7425935 | 0.3469158 | 0.0025323 | 0.0005161 | 0.0017391 | 9 |
| Gekkota | Carphodactylidae | 0.2557955 | 0.0661315 | 0.0031998 | 0.0009238 | 0.0015782 | 33 |
| Iguania | Iguania | 0.5531714 | 0.9397174 | 0.0032695 | 0.0009383 | 0.0082645 | 10 |
| Gekkota | Pygopodidae | 0.2093611 | 0.8305786 | 0.0033260 | 0.0008897 | 0.0004932 | 50 |
| Lacertoidea | Gymnopthalmidae | 0.2445580 | 0.6631081 | 0.0046525 | 0.0021050 | 0.0492061 | 13 |
| Lacertoidea | Gymnophthalmidae | 0.3604764 | 0.8755168 | 0.0055913 | 0.0009633 | 0.0000003 | 59 |
| Cordyliformes | Cordylidae | 0.6859321 | 0.3313581 | 0.0058461 | 0.0009751 | 0.0000245 | 17 |
| Iguania | Dactyloidae | 0.2471125 | 0.8302993 | 0.0061536 | 0.0030946 | 0.0819681 | 10 |
| Lacertoidea | Lacertidae | 0.5486795 | 0.7781358 | 0.0092308 | 0.0015872 | 0.0000040 | 28 |
| Iguania | Phrynosomatidae | 0.0596610 | -2.7310965 | 0.0334125 | 0.0314752 | 0.4809980 | 3 |

##### Investigating the 'Elongation Index'

Other studies have used different composite measures to accomplish the same task, notably the `elongation index` (EI) used by Title et al. (2024). In simplest terms, the elongation index is the body length divided by the body width. More specifically, this trait was calculated by imagining the animal as an idealized cylinder where length is SVL and volume is the mass, as these were traits readily available from SquamBase (Meiri 202X).

$$\text{Assuming} \quad mass = \pi r^2 SVL \quad \text{then} \quad EI = \frac{SVL}{2(\frac{mass}{\pi SVL})}$$

$$\text{Given that} \quad BodyWidth = 2(\frac{mass}{\pi SVL}) \quad \text{then} \quad EI = \frac{SVL}{BodyWidth}$$

.

**Unfortunately**, this approach was inappropriate for two reasons.

1. Length and mass data derived from SquamBase are typically maximum *recorded* values, and do not necessarily correspond to the same individual. This strongly biases the resulting elongation index values.
   - e.g. the highest-value EI species in Title et al. are *Hydrodynastes gigas*, *Pituophis melanoleucus*, and *Farancia abacura* (all EI scores >2700), species that are long, but not *notably* elongate for a snake.
   - expected high EI scoring species should be long and thin species such as typhlopid (e.g. *Ramphotyphlops longissimus*) and some arboreal colubrid (*Ahaetulla prasina*) snakes.
2. While the majority of snake mass is concentrated in the body, lizards apportion significant mass in limbs and tails. Across lizards, tails act as the primary axis of elongation, and in some species more than half the animal mass may be contained in the tail.
   - This systematically biases the elongation index towards higher values in snakes and lower values in lizards.
   - The *elongation index* of Title et al. is more an index of body elongation not absolute elongation.

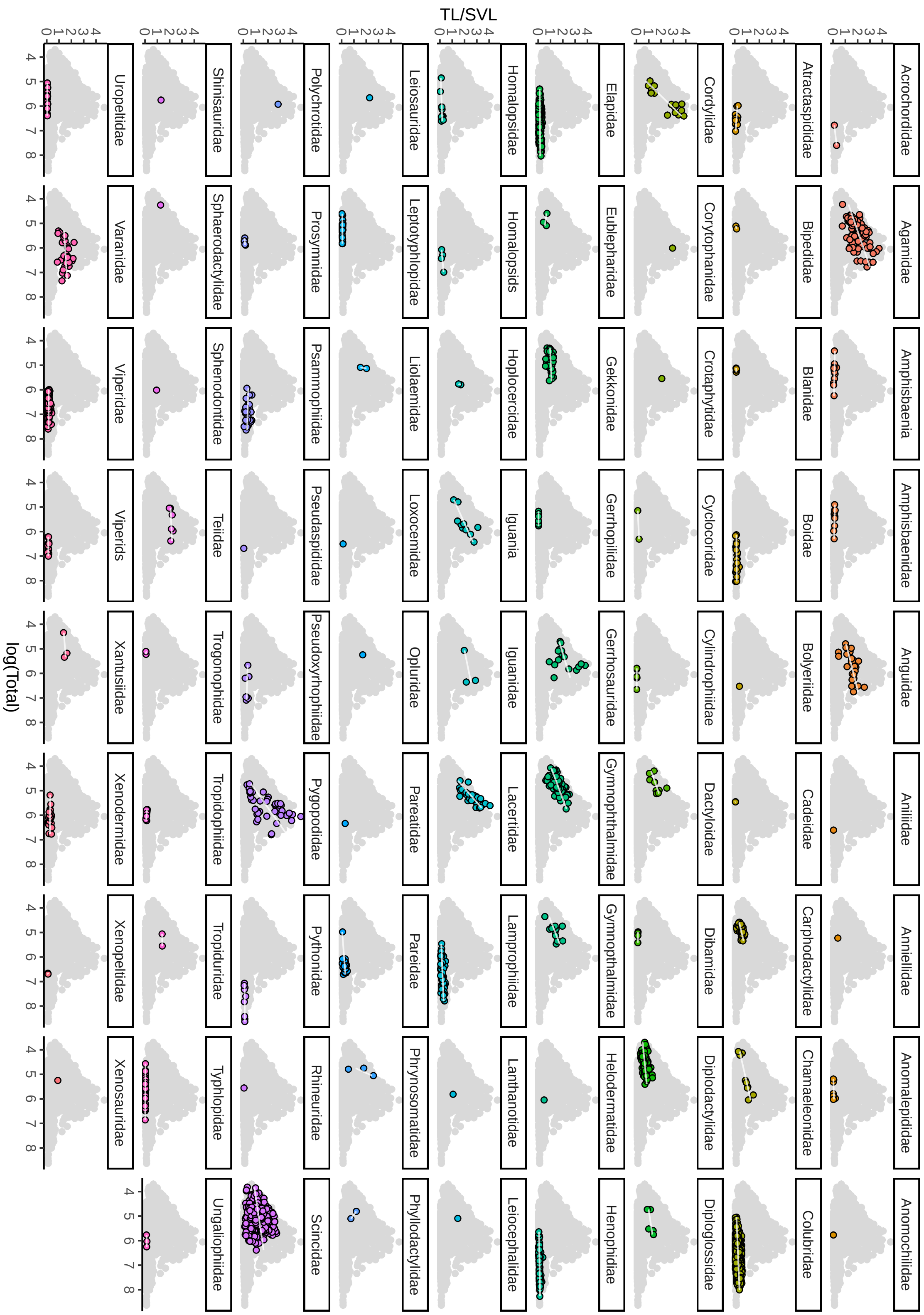

**Figure S6.** Trends in tail length to snout-vent length as a function of total length across squamate families. Where sample sizes permit, we fit linear models to family trends.

##### Modelling Tail-to-Body Evolution

To better understand the tempo and mode of elongation through tail evolution we undertook a trait modelling exercise. We started by grafting our dated pygopod tree onto the squamate topology of Title et al. (2024) and we retained squamate taxa that were available in both the tree and morphological dataset (TL/SVL). Using these data we fit a series of models ranging from an unbiased random walk to rate-variable and trended. The most basic model—Brownian Motion—allows phenotypes to evolve through incremental random change, but at deep timescales and when traits may show evolutionary rate heterogeneity this model is expected to perform poorly. The most complex—BayesTraits V4's fabric model (Pagel et al. 2022)—also follows a process of gradual change, but allows rates to shift at discrete points (changes in evolvability, $v$) on the tree and allows traits to change rapidly along individual branches through directed evolution (trends, $\beta$). We fit these models and others (Early Burst, single-peak Ornstein-Uhlenbeck) in BayesTraits V4 and estimated marginal likelihoods using a stepping-stone sampler to compare model fit.

We started by generating a BayesTraits input file for the size-corrected tail length (TL/SVL) `Data/BayesTraits/Traits/...`. We used custom functions in the `Scripts/VarRates_Functions.R` script to generate control files which include priors on evolutionary rate and ancestral trait based on the fit of a Brownian Motion model in `phytools`. See `Data/BayesTraits/Control_Files`. We also used generated a shell script to fit the models in parallel, see `Data/BayesTraits/Shells/SHELL_ALL.sh`. We fit two independent runs of each model to the data and summarized the model fits using `Scripts/01_BayesTraits_ModelComparison.R`. After identifying that the *fabric* model is the best fit to the data, we inspected the log files for stationarity (ESS > 200) and processed the *fabric* outputs to extract ancestral state values using the script `Scripts/02_Process_BayesTraits_Output_SingleChain.R`. These data were stored in `Data/Ancestral_Trait_Estimates.RData`.

Table S5: Table of comparative model fitting results. The fabric model (bold) is favored over competing models.

| model | run | marginal.likelihood |
|---|---|---|
| BM | 1 | -1370.610 |
| BM | 2 | -1370.583 |
| EB | 1 | -1308.043 |
| EB | 2 | -1308.611 |
| **fabric** | **1** | **1232.583** |
| **fabric** | **2** | **1246.020** |
| OU | 1 | -1180.268 |
| OU | 2 | -1180.628 |

To visualize changes in evolutionary rate and directed morphological changes as inferred under the *fabric* model, we used `Scripts/03_BayesTraits_Results.R`. This leans heavily on custom functions contained in `Scripts/fabric_Functions.R` to plot *fabric* outputs.

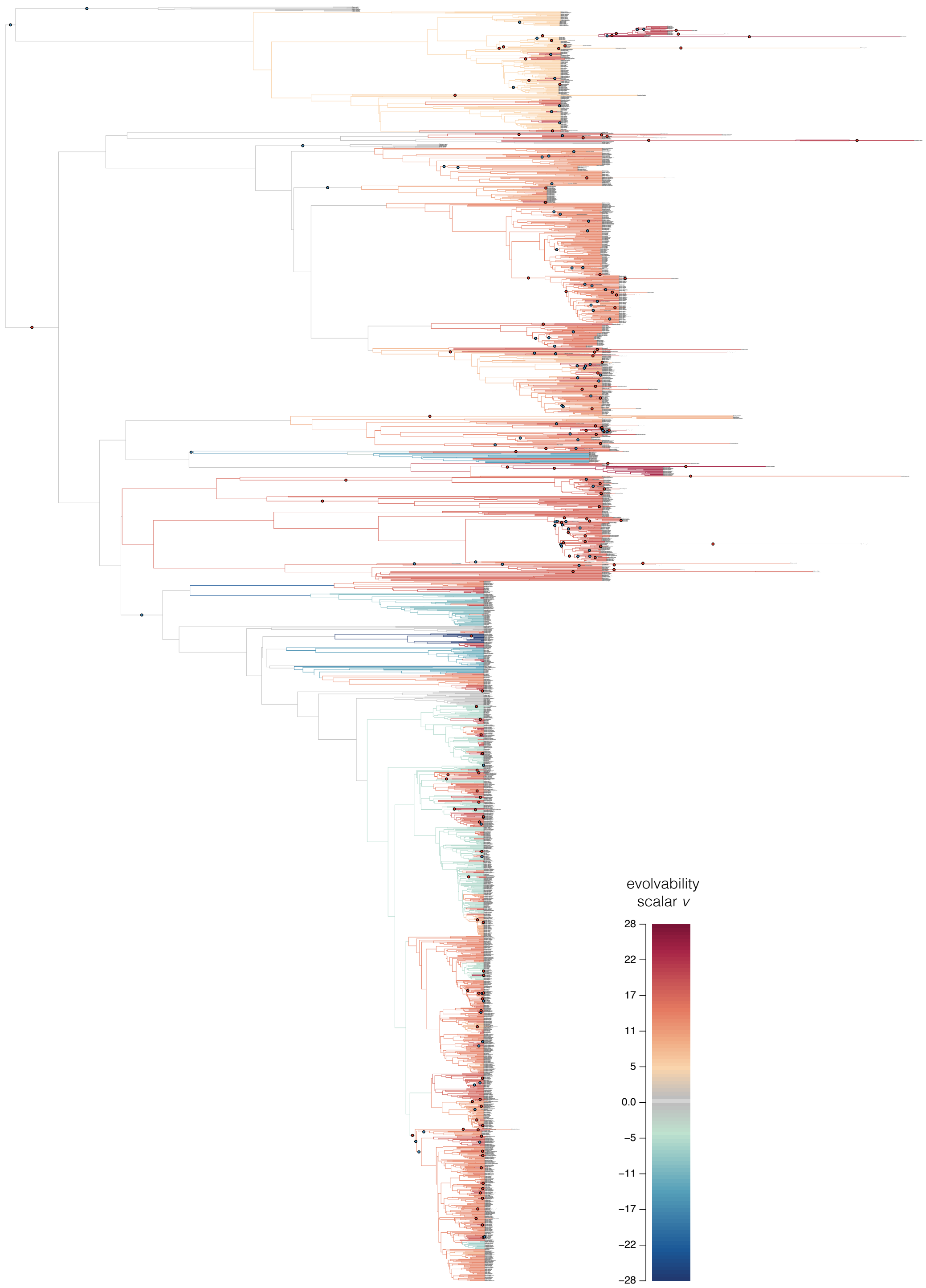

**Figure S7.** Visualization of BayesTraits fabric model fit result. Branches are colored according to evolvability scalar $v$, with warmer colors indicating faster evolutionary rates, and cooler colors indicating slower rates. Branch lengths have been scaled to reflect the phenotypic directional trend scalars $\beta$, with shortened branches indicating shifts towards shorter tail-to-body ratios, and extended branches indicating longer tail-to-body ratios. To better identify the placement of these $\beta$ shifts, we have plotted colored circles along the shifted edges, with blue circles indicating decreases and red circles indicating increases in tail-to-body ratios.

### Dietary Sampling

#### Database Assembly

To investigate patterns of diet breadth evolution across squamates we built on existing datasets (Title et al. 2024; Cavalcanti et al. 2025) by collecting data from the literature. Our dataset includes diet records for more than 90,000 individuals across 1,738 species and 63 squamate families. This new dataset significantly expands on efforts elsewhere by incorporating nearly 200 additional species, primarily lizards, including many limb-reduced species (e.g. *Dibamus*) and those with dietary items not previously captured (e.g. seaweed eating *Amblyrhynchus cristatus*, and frugivorous lizards like *Conolophus pallidus* and *Varanus mabitang*).

The dietary dataset of Title et al. (2024) (TEA24) is the largest current squamate database (>74k individuals), and while it covers all squamates, it primarily focuses on snakes. For consistency, and to harness the analytical framework presented in TEA24, we chose to add to this dataset rather than generate a new one. The second largest dataset, collated by Cavalcanti et al. (2025) (CEA25) accounts for >14k individuals and exclusively focuses on squamate lizards. To harmonize the two datasets, we recoded the dietary categories of CEA25 following those of TEA24. Those recodings are documented in *Diet/Cavalcanti_Diet_Data/Cavalcanti_ReCategorization.csv* and largely correspond to combining fine invertebrate categories into increasingly broad ones (e.g. insect_egg, insect_larvae, insect_pupae –> insect_larvae_pupae_and_eggs). We contribute an additional 63 species not covered in either TEA24 or CEA25 and provide references and DOIs for all entries (column `source_reference`). For TEA24 and CEA25 we refer readers to those publications. Unfortunately, TEA24 does not include references to primary literature for any of the 1560 species included. CEA25 includes (where available) author names, year, and journal title, and so we include this information in a separate column titled `source`. Further, dietary studies rarely provide raw data presented on a per-specimen basis. Instead, dietary information are typically presented as dietary item frequencies or proportions from a pool of individuals, and so while unsatisfying, we are unable to provide specimen-level dietary metadata.

To process and analyze the data we followed the methodology of Title et al. (2024) in standardizing diet proportions to make lizards and snakes comparable. We collected this information ("full stomach equivalents"—prey proportion x number of sampled individuals) into a table (`Diet/Squamata_Diets_Families.csv`). We then followed the analytical framework of Grundler & Rabosky (2021). Briefly, this framework calculates a phylogenetically informed estimate of dietary niche for each species, accounting for sample size variation. It accomplishes this by discretizing the multivariate diet space (32 dimensions) into K niche states (we used K = 20, 50, 100, 1000), then uses an MCMC sampler to generate posterior distributions of dietary preference across those diet variables for each taxon. Specifications for the MCMC sampler are provided in `Diet/01_mcmc-diet.R`. The result is four mcmc chains (`mcmc-...out`), one for each value of K. Results were averaged across all four chains and summarised to generate phylogenetically-smoothed dietary proportions (`Diet/diet-proportions-phylo.csv`) using the `Diet/02_make-diet-proportions.R` script.

#### Dietary Breadth Comparisons

Further dietary data exploration and analysis is documented in `Diet/Plotting_Diet_PCA2.R`. We started exploring the data by estimating dietary breadth for each species using the Inverse Simpson Index in `vegan`. We split species into one of four morphological categories (limbed lizard, limbless lizard, snake, pygopod) and plotted the distribution of dietary breadth across each group, including the group mean. For the sake of visualizing this diet space (32 dimensions) we used principle components analysis for dimensionality reduction (`prcomp`).

To compare dietary disparity among squamate families we calculated the mean pairwise distance in euclidean space among all species in each family using `dispRity` (Table S6). We then plotted mean pairwise distance against mean dietary breadth on a per-family basis to identify groups that show unique combinations broad/narrow diets and high/low dietary overlap among species within those families. This highlighted some families with remarkably conserved diets (Uropeltidae), others with diverse diets that are shared across species (Diploglossidae), and some with narrow diets that are varied among species (Colubridae, Pygopodidae, Varanidae). To quantify differences in the magnitude of dietary breadth among families, we undertook a permutation exercise comparing families in a pairwise fashion (`Scripts/05_DietaryComparison.R`). We started by translating our species-level dietary data into a distance matrix using *vegdist* in the package `vegan`, applying the Bray-Curtis method because our dietary data are proportions. We compared all combinations of families with $\geq 5$ species (n = 31 families), shuffling species between families in 999 permutations. We assessed significance through a two-sided test after correcting the p-values for multiple comparisons using the Benjamin & Hochberg (BH) method. To visualize the resulting $31^2$ comparisons we plotted results as a heatmap (Fig.S8).

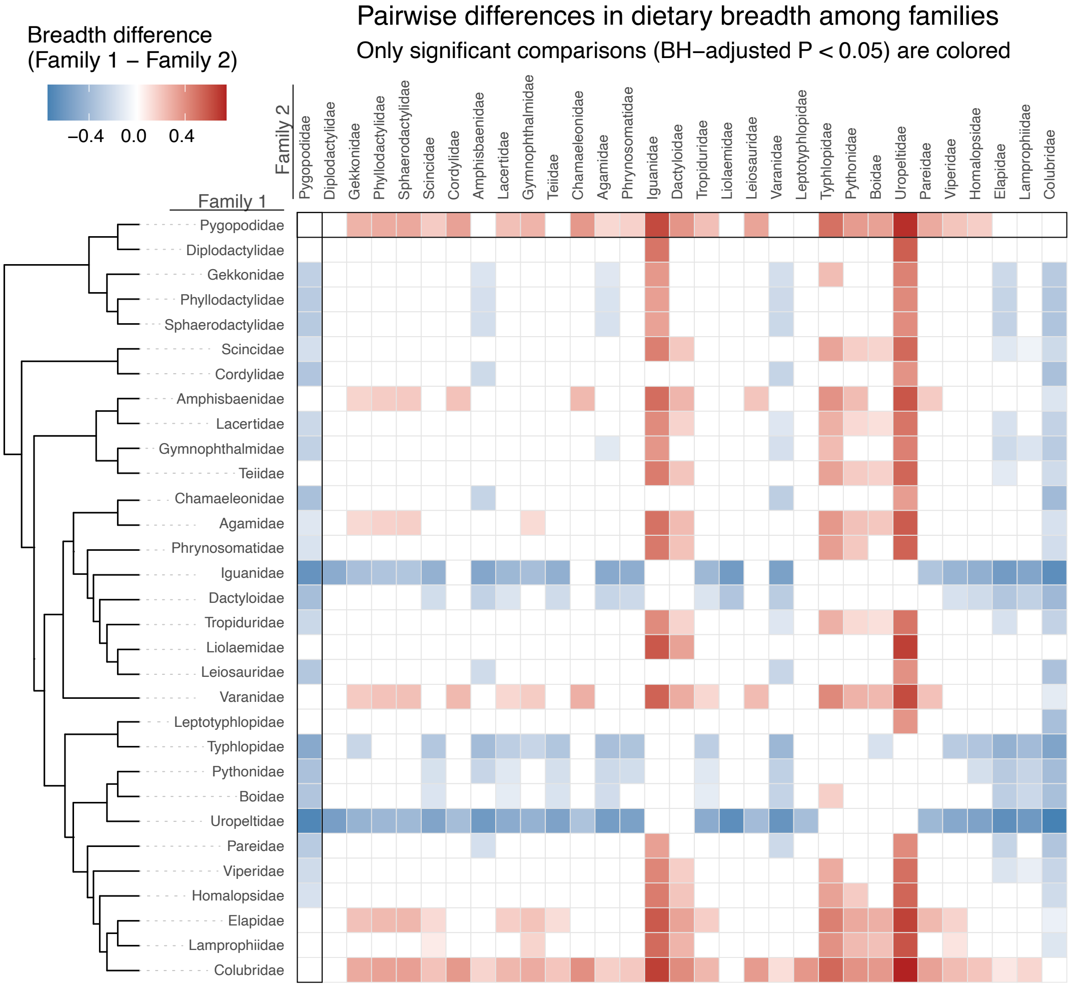

**Figure S8.** Heatmap depicting pairwise comparisons of dietary breadth between squamate families. We tested for differences in dietary breadth (as mean pairwise distance) between all pairs of families with $\geq 5$ species in our database. Non-significant results (no difference in dietary breadth) are noted by blank squares, colored squares indicate significant differences in breadth. Colors indicate the direction of difference: red indicates situations where breadth is greater in Family 1 (listed on left side of heatmap) and blue indicates situations where breadth is greater in Family 2 (listed at the top of heatmap). For example, in the comparison of Gekkonidae and Pygopodidae, the result is significant (colored) and the difference is such that Pygopodidae show a greater breadth. Worth highlighting are the trends in a small number of families: pygopod geckos, varanid lizards, and colubrid snakes show higher dietary breadths than most other squamate families. Iguanid lizards (largely herbivorous), uropeltid snakes (worm eaters), and typhlopid snakes (ants, termites, and their larvae) show much narrower dietary breadths than most other squamate families. Tree at left illustrations the phylogenetic relationships among squamate families included.

##### Dietary Breadth Simulations

To determine where these patterns in dietary breadth and variety may differ from null evolutionary expectations we estimated evolutionary rates for each dietary class across the squamate tree. To do this we fit separate Bounded Brownian Motion models in `phytools`, to each dietary category and extracted evolutionary rates (sigma) and ancestral values (theta). Using the empirically estimated values, we simulated 500 datasets with traits bounded between 0 and 1, normalized each species' dietary proportions to sum to 1, and calculated mean pairwise distance again to determine a null distribution. We then plotted each family's empirically estimated pairwise distance against the null distribution for that family to identify those that deviate from expectations (blue greater than, red less than).

Table S6: Mean pairwise distance (pd) in dietary space within each squamate family represented by >= 3 species (no.species).

| family | pd | pd.variance | no.species |
|---|---|---|---|
| Varanidae | 8.8506390 | 6.0206613 | 20 |
| Colubridae | 8.4508702 | 11.1745921 | 541 |
| Pygopodidae | 8.2776523 | 15.4526029 | 19 |
| Liolaemidae | 8.1606847 | 4.0998836 | 7 |
| Homalopsidae | 8.0575215 | 12.3966767 | 16 |
| Agamidae | 7.5232085 | 5.7869320 | 28 |
| Elapidae | 7.3337079 | 12.3546300 | 143 |
| Teiidae | 7.2069863 | 5.1095975 | 32 |
| Phrynosomatidae | 6.9227985 | 2.9608273 | 14 |
| Tropidophiidae | 6.8164902 | 14.8362955 | 4 |
| Viperidae | 6.6857410 | 8.1458534 | 109 |
| Scincidae | 6.6730315 | 4.6040518 | 68 |
| Amphisbaenidae | 6.5851825 | 4.4579439 | 13 |
| Leptotyphlopidae | 6.4858086 | 10.0050131 | 5 |
| Lacertidae | 6.4159123 | 3.9287277 | 32 |
| Tropiduridae | 6.3678174 | 3.9632569 | 31 |
| Diplodactylidae | 6.3156172 | 9.7277059 | 8 |
| Lamprophiidae | 6.2908508 | 8.5160233 | 104 |
| Gekkonidae | 6.2302060 | 3.8922042 | 16 |
| Sphaerodactylidae | 6.0067505 | 2.7442200 | 11 |
| Gymnophthalmidae | 5.7327889 | 4.3620667 | 30 |
| Phyllodactylidae | 5.6597963 | 2.8226153 | 12 |
| Cordylidae | 4.9332098 | 2.2477797 | 6 |
| Chamaeleonidae | 4.9288482 | 1.0549050 | 5 |
| Pythonidae | 4.7826496 | 5.0065927 | 18 |
| Dactyloidae | 4.7691896 | 5.1573692 | 24 |
| Leiosauridae | 4.7262750 | 6.8865725 | 6 |
| Boidae | 4.6970524 | 7.1574750 | 35 |
| Pareidae | 4.6502489 | 10.9134646 | 8 |
| Iguanidae | 4.3961330 | 6.6422631 | 8 |
| Typhlopidae | 4.2748841 | 18.4195055 | 21 |
| Diploglossidae | 3.1931577 | 1.3573430 | 3 |
| Carphodactylidae | 3.0458679 | 2.2017231 | 3 |
| Uropeltidae | 0.5937605 | 0.1704235 | 5 |

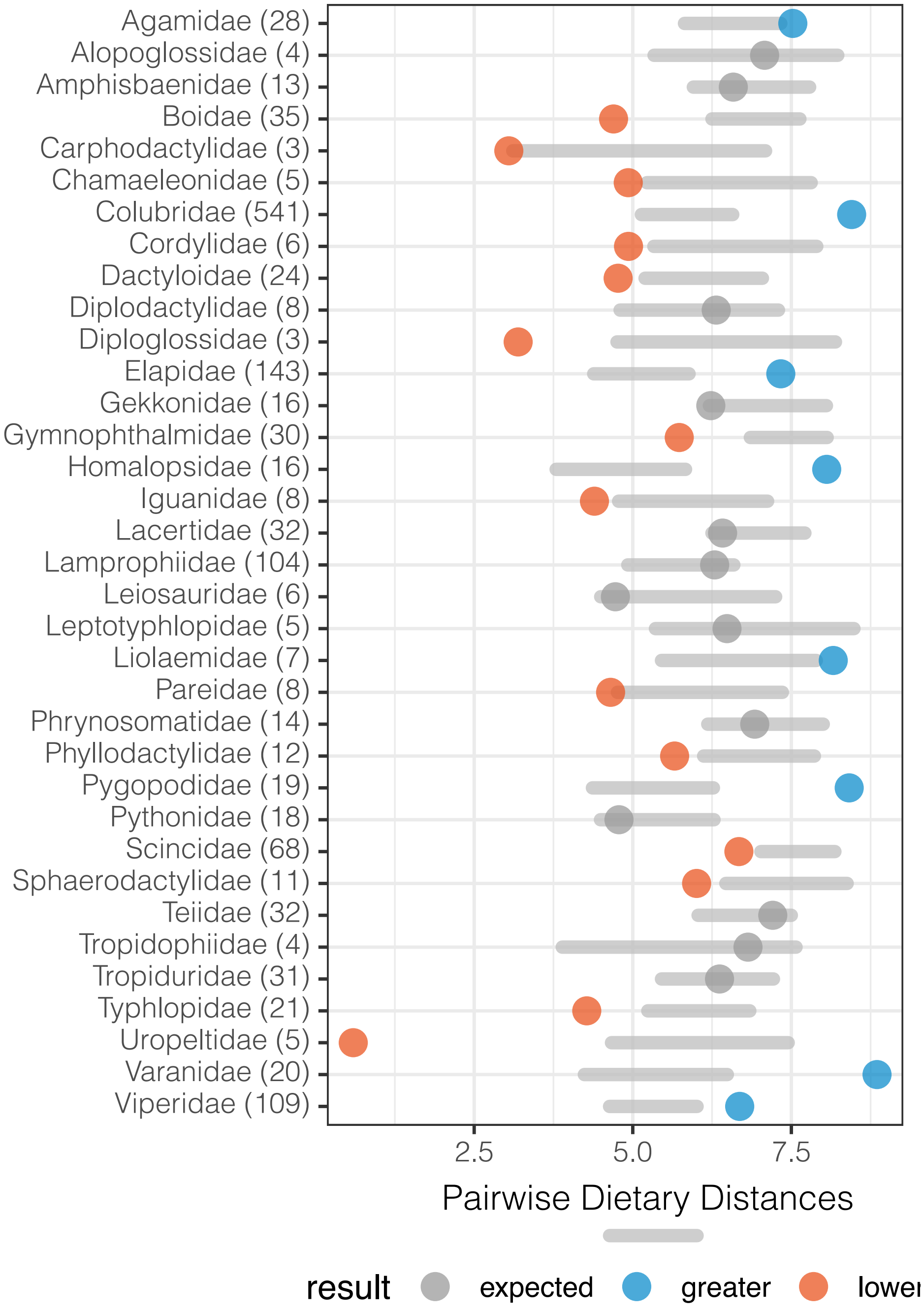

**Figure S9.** Comparison of observed dietary pairwise disparity against a null phylogenetic diet evolutionary model indicates squamate families that adhere (grey) to dietary expectations and those that deviate from expectations (greater—blue—show greater pairwise disparity; lower—red—show less pairwise disparity than expected). Families are ordered alphabetically, and number in parentheses indicates the number of sampled species.
